# Co-occurrence of Yersinia pestis and other zoonoses during European prehistory

**DOI:** 10.64898/2026.02.20.707102

**Authors:** Toni de-Dios, Biancamaria Bonucci, Jess Emma Thompson, Sofia Panella, Remi Barbieri, Marcel Keller, Tina Saupe, Sara Bernardini, Stefania Sasso, Madeleine Thirolle, Anu Solnik, Helja Kabral, Biancamaria Aranguren, Montserrat Sanz, Joan Daura, Carles Lalueza-Fox, Toomas Kivisild, Mait Metspalu, Meriam Guellil, Mary Anne Tafuri, John Robb, Christiana Lyn Scheib

## Abstract

The infection of humans by the causal agent of Plague, *Yersinia pestis*, has been attested as far back as 5,500 BP. Although the specific patho-mechanism and ultimate origin of the disease caused by these prehistoric genomes remains unclear, the bacterium spread through Europe, likely during the Late Neolithic to Bronze Age (LNBA). In this study, we analysed 9 genomic samples originating from 8 different human individuals dating to around 4950 cal BP from the site of *Grotta della Spinosa,* Tuscany, Italy. Metagenomic screening of these samples reveals one individual (GSP013) to be co-infected by *Yersinia pestis, Erysipelothrix rhusiopathiae,* and Hepatitis B virus (HBV). At least three further individuals from the site were infected with HBV, indicating its wider presence within the community. The phylogenetic placement of *Y. pestis* in GSP013 shows that this strain is closely related to the earliest LNBA Caucasus genomes of the bacterium, basal to later European diversity. This represents the earliest evidence of *Y. pestis* in the Italian peninsula (and Southern Europe more widely) to date, predating previously discovered genomes by at least 200 years. Furthermore, we retrieved 60 newly reported ancient genomes of *Erysipelothrix rhusiopathiae* and *Erysipelothrix tonsillarum* from animals and humans, dating back from 8,300 BP to 100 BP. Of these new genomes, 15 of which stem from individuals known to be infected by *Y. pestis*. This contributes to our understanding of *Y. pestis* transmission in prehistoric Europe and possible reservoirs, and offers insights into disease dynamics in communities during the 3rd millennium BCE.

## Introduction

The aetiological agent of plague, *Yersinia pestis*, is a deadly zoonotic enterobacterium. Environmental persistence is sustained through a cycle involving wild rodents, which serve as the primary hosts, and their fleas. Humans are infected only incidentally through cross-species spillover events.^1,2^. The study of this bacterium has been paradigmatic from the perspective of palaeogenomic techniques, which have been used to confirm *Y. pestis* as the culprit of the Justinianic Plague (First Plague Pandemic)^3^, and the so-called ‘Black Death’ (AD 1346–1353) and following outbreaks summarized as the Second Plague Pandemic^4^. Thanks to the development of next-generation sequencing (NGS) techniques, genetic studies have helped to shed light on the emergence, expansion, and adaptation of *Y. pestis* throughout human history^3,5^. This is the case for the first^6,7^ and second^8–10^ plague pandemics, and to a lesser extent, for the third plague pandemic^11^. Palaeogenomic studies have revealed at least 3 previously non-documented phylogenetic lineages of prehistoric *Y. pestis*: the Late Neolithic–Bronze Age *ymt* positive (LNBA+; precursor of modern diversity)^12^, the LNBA *ymt* negative (LNBA-)^13^, and the pre-LNBA clade (also missing *ymt*)^14–16^. The *ymt* gene is necessary for efficient rodent flea-mediated transmission of the bacterium^17^. The genomes of these older cases show a clear stepwise accumulation of mutations and gene pseudogenisation (for example, in *ureD*, *PDE-2*, *flhD*) that eventually enabled the bacterium’s adaptation to arthropod vectors and increased its virulence in mammalian hosts as we know it today^12,13^. Accordingly, the gene profiles of these prehistoric genomes suggest that plague transmission to humans during the LNBA was unlikely to have relied on the classical rodent–rodent flea–human cycle, as is predominantly known to be the case for the third plague pandemic and modern outbreaks^18,19^.^.^ Moreover, the role of rodent fleas in plague transmission during the second pandemic remains debated, with bubonic plague likely transmitted primarily via human ectoparasites including human fleas and body lice^20,21^. For LNBA-genomes, the exact transmission route remains unclear.

The prehistoric lineages have provided estimates from the most recent common ancestor of *Y. pestis* diversity, dating to between 11,300 and 7,300 before present (BP)^22^. Yet the picture of the emergence and spread of prehistoric *Y. pestis* genomes remains puzzling, with an unclear geographical origin somewhere in Eurasia. Current palaeo-pathogenomics studies place the deepest lineages of *Y. pestis* at 5,600 to 5,000 BP around the Lake Baikal area^22^, with a derived strain appearing in what is now Latvia (RV2039; 5,300–5,000 BP)^16^. These prehistoric, seemingly extinct, lineages are distinct from the later lineages of the more recent historic pandemics. Indeed, prehistoric *Y. pestis* lineages display far less clonal expansion^23^ and fall on multiple branches, which are often represented by single genomes. Despite lacking part of the repertoire of virulence genes present in modern genomes, archaeological data suggest that prehistoric *Y. pestis* genomes were virulent^22^, caused a sustained burden on the affected populations^24^ and has been debated as having contributed to major population shifts^14^.

Modern parallels show that domestic animals can act as bridge hosts, contracting plague through interaction with wild rodents or other mammals^25–27^. Those domestic animals have been known to then transmit the bacterium to humans without the need for flea vectors^28–33^. Indeed, recent studies have reported the presence of *Y. pestis* in a Neolithic dog in Sweden^15^, and in a domesticated Bronze Age sheep from Central Asia^34^, this latter example is genetically close to human LNBA genomes. Although minimal thus far, this evidence could support a model in which animal husbandry played a role in local plague transmission during prehistory, creating new epidemiological scenarios for spillover events from wildlife and domestic animals to humans or vice versa^23^. This is particularly true for dogs^32^, which, much like rodents, can be carriers of the bacterium without suffering severe adverse effects^35^.

Another zoonotic pathogen capable of infecting humans is *Erysipelothrix rhusiopathiae*, which has been sparsely reported in ancient genomic datasets^36,37^. *E. rhusiopathiae* is an opportunistic, gram-positive, non-spore forming, facultative intracellular pathogen with a wide range of hosts^38^. *E. rhusiopathiae* is commonly associated with livestock, primarily infecting pigs’ tonsils, causing asymptomatic infection and swine erysipela^38^ (skin lesions, necrosis, and death), but can also affect other domestic and wild ungulates^39,40^ and occasionally causes epizootic outbreaks in both wildlife and domestic animals^40–42^. Other evolutionarily distant hosts can be infected by the bacterium, with outbreaks affecting animals such as dogs^43^, marine mammals^44^, birds^45^, and fishes^46^, among others. Humans can be infected with *E. rhusiopathiae*, through close contact with infected animals, contaminated soil or water, with farmers, butchers, fishermen, and other professionals at risk^47^. The bacterium can remain infectious in soil for up to 35 days^48^. The typical manifestation of *E. rhusiopathiae* in humans is *Erysipeloid of Rosenbach,* a skin condition causing skin lesions^38^. The symptoms in humans are similar to those of pigs^38^, with a systematic review of 62 case studies of *E. rhusiopathiae* infections from 2000 to 2020 revealing that the main clinical manifestations of the disease in humans are: fever (56.45% of cases), pain (32.45%), localised skin lesions (‘erysipeloids’; 29.03%), and endocarditis (29.03%)^49^. Other severe manifestations of the disease include sepsis (12.9%), abscesses (6.45%), generalised skin lesions (4.83%), and meningitis (3.22%)^49^. Most cases of septicemia are linked to endocarditis, with a mortality rate of 38%, one of the highest for bacterial endocarditis despite the fact that, proportionally, it is a rare cause of bacterial endocarditis^47^. Genomically, this species presents low levels of clonality and a high degree of recombination, with 3 generalistic main clades infecting a wide range of hosts that do not group by host or geographical origin^50^. Nowadays, infections of humans by *E. rhusiopathiae* are rare, however, we expect that they could have occurred in higher frequencies in the past due to differences in living conditions and husbandry, but its presence in the past remains mostly unstudied.

In this study, we have retrieved genetic data for 8 human individuals from the Copper Age Central Italian site of *Grotta della Spinosa* [See Methods section: “*Archaeological context of the Grotta della Spinosa, Italy”*, Figure 1, Supplementary Figures S1-S4]. One individual (GSP013) presented evidence of infection by multiple pathogens, including a LNBA- *Y. pestis* strain and *E. rhusiopathiae,* as well as Hepatitis B virus. We also present evidence of *Erysipelothrix spp*. in ancient animals and humans, and co-occurrence of *Y. pestis* and *Erysipelothrix rhusiopathiae* and *E. tonsillarum* or *Salmonella enterica* and *E. rhusiopathiae* in multiple published ancient individuals. Our data provides a new geographical and temporal boundary for LNBA- *Y. pestis* in Western Europe, and substantially increases the number of ancient *Erypselothrix* genomes available. This could help to better understand how *Y. pestis* spread throughout prehistoric Europe, and how the disease caused by *Erypselothrix* spp. affected ancient human communities as well as its relevance in the context of zoonotic disease transmission.

**Figure 1.**
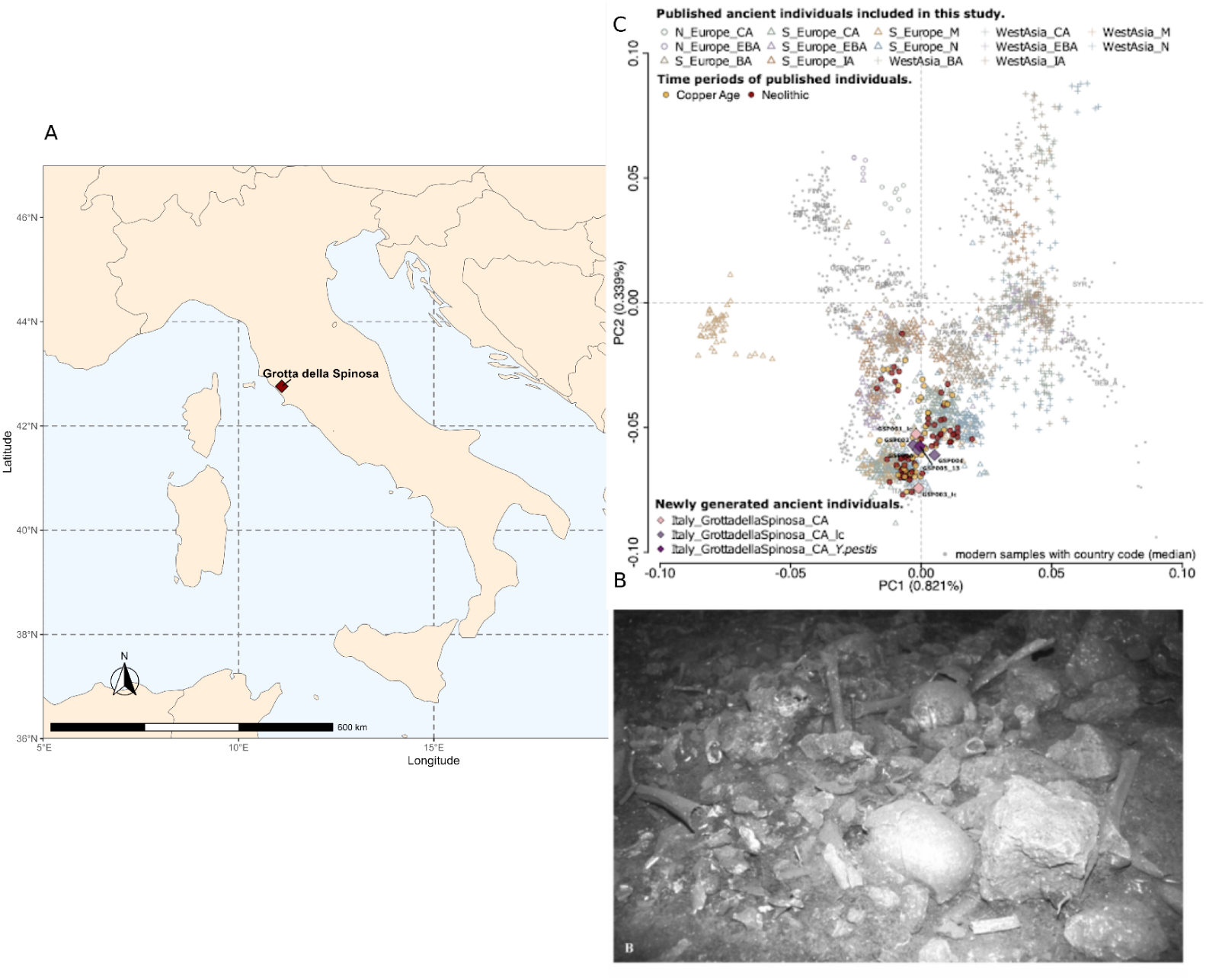
Site location and Principal Component Analysis of the human individuals. (A) Location of the *Grotta della Spinosa* site within Italy. (B) Photograph of the commingled skeletal remains found at the site. Permission to reuse the image granted by Dr. Biancamaria Aranguren. (C) PCA of the newly generated and published ancient human individuals projected onto the genetic variation of present-day Eurasian populations. lc - low coverage. _N - Neolithic, _CA - Copper Age, _(E)BA - (Early) Bronze Age, _IA - Iron Age, _M - Mesolithic

## Results

### DNA Screening and individual selection

To investigate the possible presence of pathogens in our samples, as well as the genetic ancestries and biological relatedness of the human individuals buried at *Grotta della Spinosa* (GSP), we initially selected teeth (n = 3) and skeletal elements (auditory ossicle = 1, petrous portion of temporal bone = 6) from what we assumed were nine different individuals: seven adults (adult (18+ years old (yo)) = 4, young-mid adult (25–35 yo) = 1, middle adult (35–45 yo) = 1, mature adult (45–60 yo) = 1) and two non-adults (adolescent 12–18 yo) [Supplementary Information S1: Selection of archaeological samples]. The remains were entirely disarticulated and commingled. In an effort to avoid duplicate samples we prioritised the left petrous portion of the temporal bone, which was most numerous, and we selected the same type of molars, so that could be attributed to different individuals. Teeth were preferably extracted from mandibles, which had been previously analysed for stable isotopes^51^. We extracted DNA from these skeletal remains and built double-indexed double-stranded DNA (dsDNA) libraries in the ancient DNA facility at the Institute of Genomics, University of Tartu (Estonia) [See Material and Methods for details, Supplementary Table S1A-B].

We sequenced nine skeletal samples generating a total of 176,415,256 reads [Supplementary Table S1C]. After sequencing, we used READv2^52^ to determine identical individuals in the given sample set. This analysis confirmed samples GSP005 (a petrous portion) and GSP013 (a tooth) [Supplementary Figure S4] to be identical, suggesting they originated from the same individual (referred to as GSP013 in the text), leaving eight unique individuals with a final average human coverage of 0.26× (range: 0.0016× - 0.834× , median = 0.15×) and modern human mtDNA contamination between 0.1 - 7.3% [Supplementary Table S1C].

In parallel, we deduplicated and removed low complexity sequences from the trimmed data, and used this filtered data to perform a metagenomic screening of the libraries using KrakenUniq, validating possible hits by mapping (See Methods). We observed sequences potentially associated with *Yersinia pestis* and *Erysipelothrix rhusiopathiae* in the GSP013 libraries [Supplementary Table S2, Figures S5-S6]. Additionally, we found evidence of *Hepatitis B virus (HBV)* infection in 4 of the 8 analysed individuals (GSP004, GSP011, GSP013, GSP014) [Table 1; Supplementary Figure S7-S8; Supplementary Table S2]. To increase the yield of genomic data, we performed capture of the *Y. pestis* and *Y. pseudotuberculosis* genomes, in addition to a second round of shotgun sequencing of the original library. From GSP013, we retrieved a complete *Y. pestis* genome at an average depth of 3.26× coverage, a partial *E. rhusiopathiae* genome at 0.515×, and a partial *HBV* sequence at 0.756× [Table 1; Supplementary Figure S9].

**Table 1.** Mapping statistics. GSP013 and GSP014 basic mapping statistics against *Y. pestis CO92, E. rhusiopathiae NCTC8163* (GCF_900637845) and *HBV ayw* reference genomes.

| Individual | Organism | Q1 Reads | Unique Q30 Reads | Average Depth | Breadth of Coverage |
| --- | --- | --- | --- | --- | --- |
| GSP013 | *Y. pestis* CO92 (Chromosome) | 4,635,707 | 295,257 | 3.232× | 81.14% |
| GSP013 | *Y. pestis* CO92 (*pCD1*) | 120,328 | 7,460 | 5.539× | 86.16% |
| GSP013 | *Y. pestis* CO92 (*pMT1*) | 45,999 | 3,708 | 1.932× | 60.11% |
| GSP013 | *Y. pestis* CO92 (*pPCP1*) | 39,592 | 2,502 | 14.143× | 79.12% |
| GSP013 | *E. rhusiopathiae* | 28,245 | 17,663 | 0.515× | 38.73% |
| GSP013 | Hepatitis B virus | 173 | 36 | 0.756× | 28.63% |
| GSP014 | Hepatitis B virus | 179 | 121 | 1.846× | 55.69% |
| GSP004 | Hepatitis B virus | 1 | 1 | 0.0116× | 1.16% |
| GSP011 | Hepatitis B virus | 5 | 3 | 0.0374× | 3.74% |

Finally, we assessed the chronology of the deposition of the commingled human remains within *Grotta della Spinosa*. The sample for GSP013 was obtained from a tooth extracted from a mandible in level 3 of stratigraphic layer (US) 8 [Supplementary Figure S5], from an adolescent (16–18 years) with no apparent dental pathology [See Supplementary Information S2: Paleopathological analysis of GSP013]. Bayesian analysis of published radiocarbon dates shows that the burials were deposited between the late 4th and early 3rd millennium BCE^53^, for a period of 120 to 490 years (2*σ*) [see Supplementary Figure S10]. GSP013’s mandible was excavated from quadrant G1. A previously obtained radiocarbon date from an ulna found in quadrant G1, in the same stratigraphic layer and level, places the sample at the turn of the 3rd millennium BCE: 3100–2920, 2σ (Beta-456210: 4370±30 BP, calibrated in OxCal v4.4.4 with Intcal20)^53–55^ [see Supplementary Table S3]. Despite the disarticulation and commingling of the bone deposits, the substantial number of consistent dates from this stratigraphic layer indicate that the date from the ulna in G1 can be used as a reliable representation of the horizon of individual GSP013’s lifetime, likely between 5050–4850 cal BP.

### Genetic ancestries of Grotta della Spinosa

To prepare the selected datasets for genome-wide analysis, we called the single-nucleotide polymorphisms (SNPs) on the ‘1240k’ panel^56^ using ANGSD^57^, retaining an average 156,000 SNPs per sample (approximately ranging from 1,000 to 450,000 positions, median = ∼120,000). All individuals, except for GSP011 and GSP014 (with <20,000 overlapping SNPs), were merged with publicly available datasets from West Eurasia^58–84^ using the Allen Ancient DNA Resource (AADR) data (version v62.0)^85,86^. To investigate the genetic ancestries of the human individuals from *Grotta della Spinosa*, we projected the newly generated and the publicly available ancient individuals onto the genetic variation of present-day Eurasian populations [Figure 1, Material and Methods, Supplementary Table S4]. We found that all newly generated individuals from the site fall into the so-called European Neolithic cluster, presenting a homogenous ancestral composition to be expected from a Copper Age Italian population.

### An early case of Yersinia pestis in Southern Europe

With the aim to understand the phylogenetic positioning of the *Y. pestis* strain found in individual GSP013 in relation to other prehistoric genomes of *Y. pestis*, we built a genome-wide, high-coverage SNP dataset. The final dataset consisted of 37 high quality (≥ 3 fold average depth) ancient *Y. pestis* genomes in human individuals and GSP013 (Neolithic, LNBA+ and LNBA-), mapped against *Y. pestis* strain CO92^12–14,16,23,87–89^, and with *Y. pseudotuberculosis* IP32881^90^ as root [Supplementary Table S5]. We created a Maximum Likelihood (ML) phylogenetic tree using raxml-ng^91^ and 1,042 variant sites [Figure 2A][See Methods]. Additional trees without singletons, without transitions, and with higher genotype quality thresholds, were generated to account for false positive sites produced by terminal damage or low coverage [Supplementary Figure S11]. GSP013 falls within the diversity of the LNBA-clade of *Y. pestis* (Transfer Bootstrap Expectation (TBE) value=0.9998; average tree TBE=0.9771), basal to all other genomes with the exception of the Northern Caucasus strain RK1001 (4828–4622 cal BP)^88^, and a strain from the Altai region, RISE509 (4837–4627 cal BP)^13^. This would represent the oldest currently attested evidence of *Y. pestis* LNBA-infection in Europe, predating the later Central and Eastern European genomes by around 200 years.

**Figure 2.**
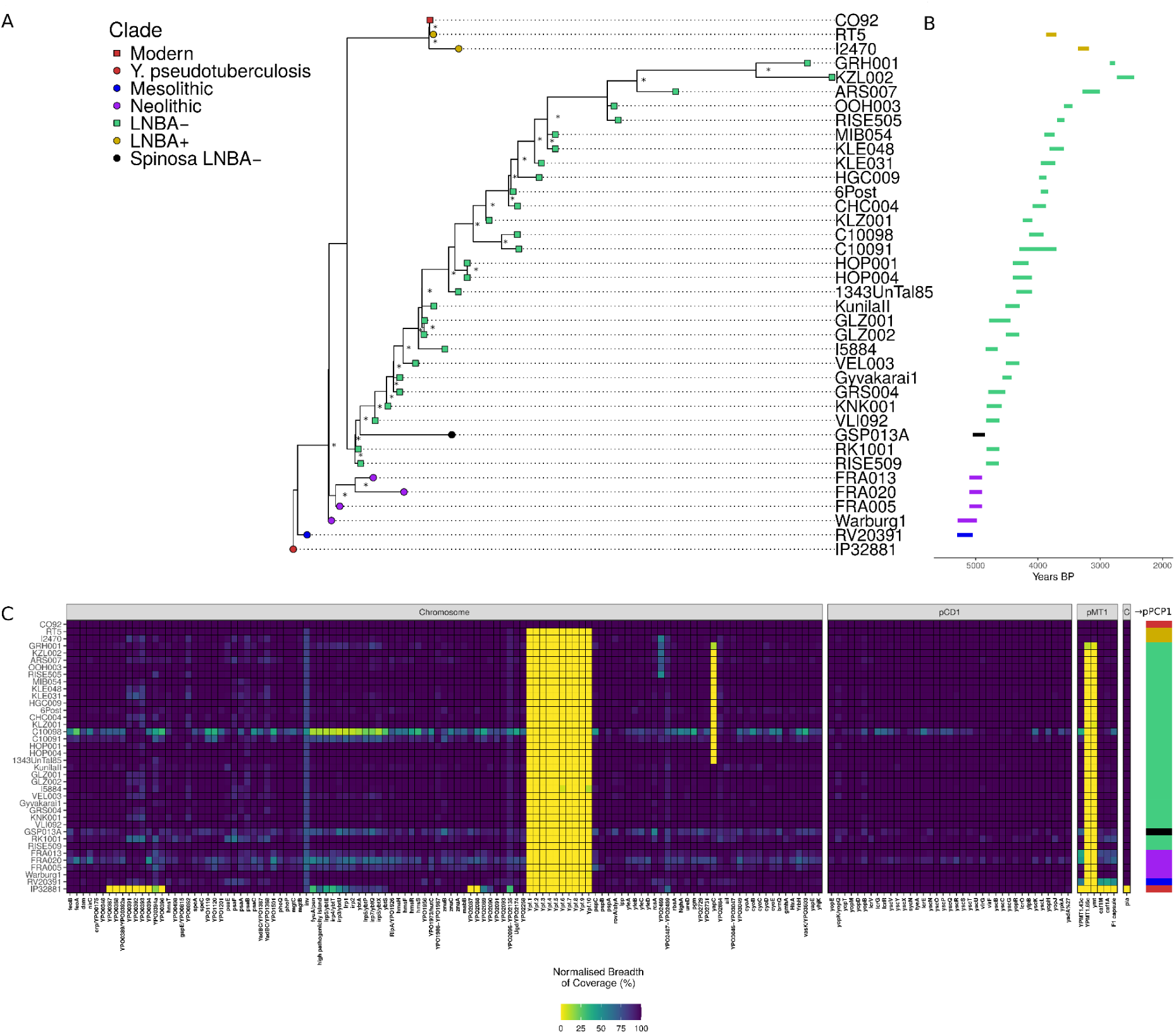
Phylogenetic relationships of GSP013. (A) Maximum Likelihood tree of GSP013 strain and 37 published ancient and modern *Y. pestis* genomes. The tree is rooted to *Y. pseudotuberculosis* strain IP32881. We have used GTR+G and 1,000 bootstraps. Nodes with a TBE bootstrap support ≥ 0.9 are marked with (*) at the base of the node. GSP013A has been colour-highlated inside clade LNBA-. (B) Dating estimates for each sample in years before present (BP). (C) Heatmap of inferred presence/absence of virulence-associated genes assessed by normalised depth-of-coverage across the *Yersinia pestis* Chromosome and plasmids pCD1, pMT1 and pPCP1 with the colour bar denoting the Clade.

**Figure 3.**
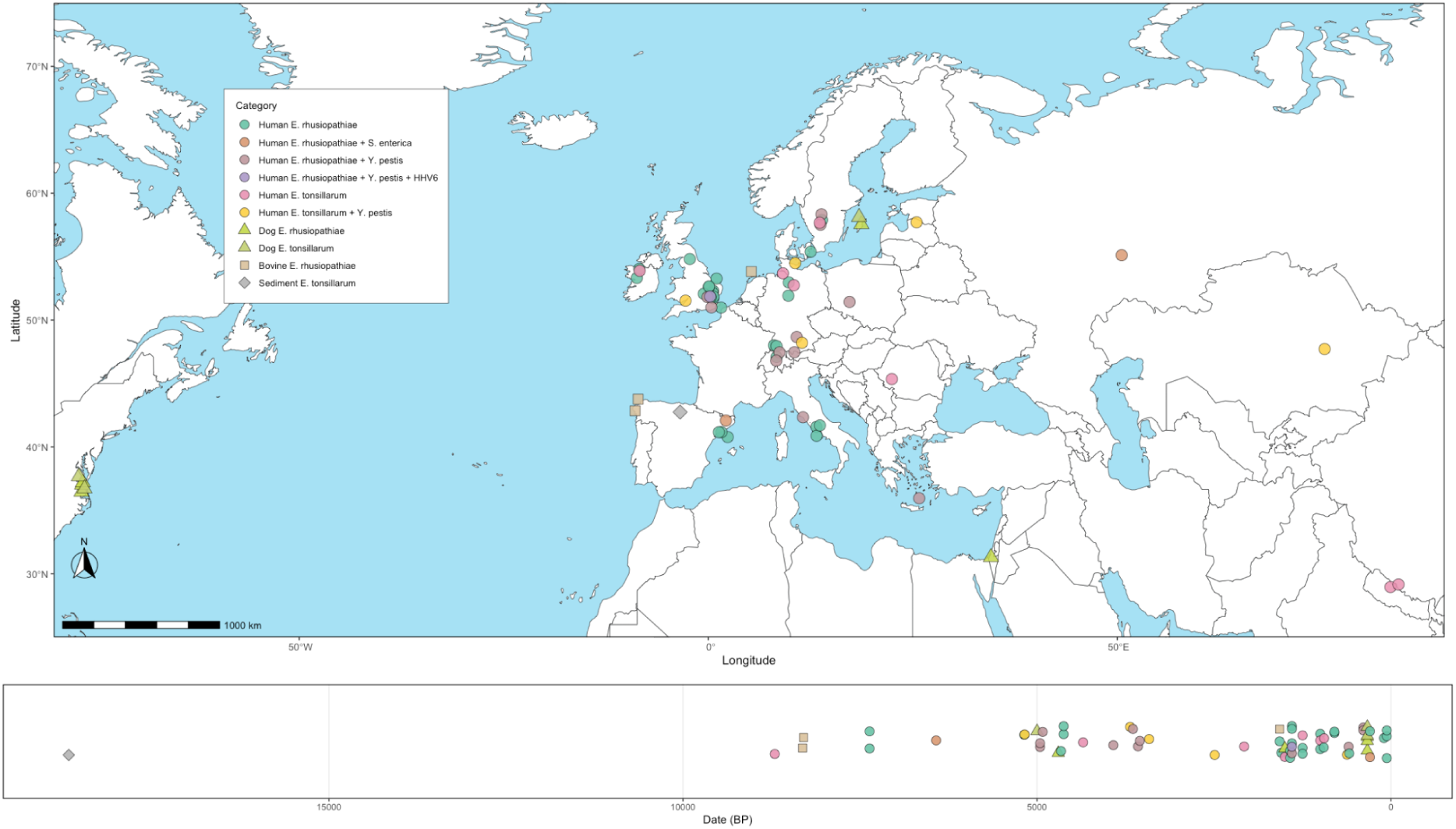
Geographic and temporal distribution of *Erysipelothrix* spp. genomes reported in this study. Map with the geographical origin of the retrieved genomes, one dimensional transect with the estimated age of the genomes.

Next, we evaluated the presence and absence of known *Y. pestis* virulence factors in the reference genome (CO92) for the sample reported here, and compared the results with other samples from previous studies^8,13–16,23,24,88,89,92^ [Figure 2B]. We observed the absence of the filamentous prophage, which is missing in all published Neolithic and LNBA genomes, and today is mostly found in 1.ORI genomes^93^. As expected, GSP013 also lacks both the *ymt* gene, which enables *Y. pestis* to survive within the midgut of the flea^94^, crucial for flea infection in a broad range of hosts^95,96^; and YPMT1.66c, which is involved in resistance to mammalian innate immunity^97^. We also observe the absence of the *yapC* gene, which has been shown *in vitro* to mediate cell adhesion and facilitate biofilm formation^98^. We manually explored the *pla* genotype of GSP013 and we found that it exhibited the ancestral allele, which has been associated with a less efficient dissemination in mammals compared with the derived T259 allele^99^. GSP013 seemed to present functional variants of the virulence-associated genes *ureD*, *PDE-2* and *flhD,* which were manually explored, but the coverage was too low to draw any further conclusion [see Supplementary Information S3; Supplementary Figure S12].

### Erysipelothrix as an indirect marker of zoonotic contacts

The presence of *E. rhusiopathiae* in GSP013 hints at a possible contact with an infected animal close to the individual’s death. The disease has an estimated incubation time in pigs of 1 to 7 seven days in acute infections, 2 to 4 weeks in chronic manifestations of the disease, with cases manifesting between 1 and 20 years after the infection^100,101^. It is believed that the number of diagnoses with this bacteria may be underestimated due to the low effective growth of the bacterium in culture^102^ and other difficulties associated with its identification^103^. The number of published ancient genomes belonging to this bacterium is low (4 published)^36,37,104^. We screened 2,495 individuals from published ancient and historical datasets for the presence of *Erysipelothrix spp.* [see Supplementary Information SI4 and Supplementary Table S6]. Those include sites at which at least one *Y. pestis* or *S. enterica*^3,5–9,12–14,16,23,24,61,88,89,92,105–121^ infection was present, as well as ancient dogs and wolves^122–124^, ancient ovine and bovine genomes^34,125–127^. We also analyzed unpublished metagenomic library data from 6 previously published Neolithic individuals^128^. We additionally included 77 metagenomic samples from archaeological sediments to test if *Erysipelothrix spp.* can be found as an environmental contaminant in contexts associated with animals or humans^107,127,129–132^ [Table 2; Supplementary Table S6]. We mapped all collected data against a set of 12 different *Erysipelothrix spp.* genome assemblies, including *E. rhusiopathiae*^133–135^*, E. larvae*^134^*, E. tonsillarum*^134^*, E. piscisicarius*^136^*, E. anatis*^137^*, E. aquatica*^137^*, E. urinaevulpis*^137^*, E. inopinata*^138^*, E. enhydrae*^134^*, and E. amsterdamensis*^139^ [see Supplementary Table S6]. Following this initial mapping, we selected the best candidates based on the number of mapped reads and the average edit distance against the assembly. We considered as positive hits the ones that simultaneously displayed a depth of coverage above 0.01× and a breadth of 1%, while also showing decreasing edit distance values with an average edit distance below 1.5. Other factors such as terminal deamination, PMD scores, and read length distribution (for samples with UDG treatment) were taken into account [Supplementary Table S6]. A final species confirmation was accomplished by projecting with epa-ng^140^ the retrieved genome to a core genome ML phylogeny, built from 636 core genes shared between *E. tonsillarum* and *E. rhusiopathiae* [Figure S13]. In the case of outliers ADN004 and C10091, further investigation of the reads reveal the presence of *Erisipelothrix*-like sequences mapped. Those two samples were excluded from further analysis. We also discarded a *E. rhusiopathiae* genome 10× from a human sample due to the lack of metadata (15594A)^141^.

**Table. 2.** Newly reported *E. rhusiopathiae* and *E. tonsillarum* genomes. Strain name, date (BP), geographical origin, coverage, host information and bacterial presence status of the new *E. rhusiopathiae* and *E. tonsillarum* genomes found in published data.

| Sample | Species | Date (BP) | Country | Breadth of Coverage | Av. Depth | Av. Length | 5' Dam | 3' Dam | Host | Skeletal Element | Library type | UDG treatment | Library Strategy | Detected Pathogens | Publication |
| --- | --- | --- | --- | --- | --- | --- | --- | --- | --- | --- | --- | --- | --- | --- | --- |
| EMN005 | *E. tonsillarum* | 18675±435 | Spain | 24.73% | 0.2976× | 46.669 | 37.78% | 41.91% | Sediment | Sediment | dsDNA | none | Shotgun | - | 130 |
| OC1 | *E. tonsillarum* | 8,704±269 | Romania | 3.33% | 0.034× | 118.65 | 23.25% | 21.32% | Human | Petrous | dsDNA | none | Shotgun | - | 191 |
| Galic3 | *E. rhusiopathiae* | 8311±60 | Spain | 91% | 65.389× | 75.62 | 2.91% | 3.84% | Auroch | Humerus | dsDNA | full | Shotgun | - | 125 |
| Galic1 | *E. rhusiopathiae* | 8297±59 | Spain | 4.3% | 0.0443× | 68.019 | 4.76% | 4.23% | Auroch | Petrous/Humerus | dsDNA | full | Shotgun | - | 125 |
| CB13 | *E. rhusiopathiae* | 7365±55 | Spain | 92.93% | 34.076× | 48.719 | 33.48% | 33.27% | Human | Tooth | dsDNA | none | Shotgun | - | 128 |
| MUR009 | *E. rhusiopathiae* | 6425±75 | Russia | 7.11% | 0.0831× | 52.714 | 0.62% | 1.13% | Human | Tooth | dsDNA | full | Capture | *S. enterica* | 105 |
| FRA012 | *E. rhusiopathiae* | 5180 | Sweden | 91.4% | 11.726× | 58.2025 | 2.82% | 2.94% | Human | Tooth | dsDNA | full | Shotgun | - | 24 |
| RV20391 | *E. tonsillarum* | 5175±125 | Latvia | 2.6% | 0.0308× | 49.603 | 4.76% | 7.61% | Human | Tooth | dsDNA | half | Shotgun | *Y. pestis* | 16 |
| C88 | *E. tonsillarum* | ~5000 | Sweden | 89.46% | 4.465× | 56.582 | 1.00% | 1.04% | Dog | Tooth | dsDNA | full | Shotgun | - | 122 |
| FRA005 | *E. rhusiopathiae* | 4957±85 | Sweden | 14.62% | 0.1716× | 41.103 | 12.65% | 2.99% | Human | Tooth | ssDNA/dsDNA | none/full | Shotgun/Capture | *Y. pestis* | 24 |
| GSP013 | *E. rhusiopathiae* | 4920±50 | Italy | 38.73% | 0.5155× | 51.67 | 44.31% | 45.46% | Human | Tooth | dsDNA | none | Shotgun | *Y. pestis* | This study |
| C89 | *E. rhusiopathiae* | 4700±200 | Sweden | 76.78% | 1.664× | 57.856 | 0.86% | 0.87% | Dog | Tooth | dsDNA | full | Shotgun | *Y. pestis* | 122 |
| LSC005 | *E. rhusiopathiae* | 4661±136 | Italy | 91.05% | 5.675× | 43.5703 | 25.48% | 24.99% | Human | Tooth | dsDNA | none | Shotgun | - | 84 |
| LSC003 | *E. rhusiopathiae* | 4624±175 | Italy | 11.66% | 0.1253× | 46.4209 | 29.38% | 29.71% | Human | Tooth | dsDNA | none | Shotgun | - | 84 |
| LSC009 | *E. rhusiopathiae* | 4624±175 | Italy | 23.37% | 0.2729× | 44.6259 | 25.69% | 26.23% | Human | Tooth | dsDNA | none | Shotgun | - | 84 |
| FRA202 | *E. tonsillarum* | 4349 | Sweden | 10.73% | 0.1876× | 47.484 | 0.73% | 1.27% | Human | Tooth | dsDNA | full | Shotgun | - | 24 |
| HGC009 | *E. rhusiopathiae* | 3922±64 | Greece | 18.25% | 0.2323× | 44.906 | 27.21% | 11.33% | Human | Tooth | ssDNA/dsDNA | none | Capture | *Y. pestis* | 89 |
| C10091 | *E. tonsillarum* | 3685±30 | England | 10.47% | 0.1846× | 46.225 | 15.69% | 2.69% | Human | Tooth | ssDNA | none | Shotgun | *Y. pestis* | 92 |
| RISE139 | *E. rhusiopathiae* | 3645±33 | Poland | 2.33% | 0.0239× | 38.045 | 15.41% | 14.87% | Human | Tooth | dsDNA | none | Shotgun | *Y. pestis* | 13 |
| 6Post | *E. rhusiopathiae* | 3574±19 | Germany | 91.49% | 38.098× | 63.255 | 7.06% | 6.73% | Human | Tooth | dsDNA | half | Shotgun | *Y. pestis* | 88 |
| KLE031A | *E. rhusiopathiae* | 3548±34 | Germany | 27.89% | 0.3708× | 52.628 | 11.3% | 8.91% | Human | Tooth | dsDNA | full | Capture | *Y. pestis* | 23 |
| KLE048A | *E. tonsillarum* | 3417±27 | Germany | 6.45% | 0.0731× | 47.369 | 16.52% | 11.51% | Human | Tooth | dsDNA | full | Capture | *Y. pestis* | 23 |
| KZL002 | *E. tonsillarum* | 2491±33 | Kazakhstan | 5.64% | 0.0759× | 60.952 | 5.67% | 7.25% | Human | Tooth | dsDNA | half | Capture | *Y. pestis* | 23 |
| TGEZ06 | *E. rhusiopathiae* | 2400±100 | Israel | 8.01% | 0.0855× | 49.478 | 20.66% | 25.22% | Dog | Petrous | dsDNA | none | Shotgun | - | 122 |
| M63 | *E. tonsillarum* | 2075±225 | Nepal | 79.61% | 2.383× | 74.7792 | 7.51% | 7.70% | Human | Tooth | dsDNA | none | Shotgun | - | 169 |
| ELY005 | *E. rhusiopathiae* | 1574±19 | England | 1.81% | 0.0127× | 50.8587 | 3.28% | 3.26% | Human | Petrous | dsDNA | half | Shotgun | - | 142 |
| Win1 | *E. rhusiopathiae* | 1573±72 | Netherlands | 11% | 0.1196× | 56.278 | 1.63% | 1.49% | Cattle | Petrous | dsDNA | full | Shotgun | - | 126 |
| BUK005 | *E. rhusiopathiae* | 1550±19 | England | 20.97% | 0.2408× | 54.7154 | 17.63% | 1.47% | Human | Petrous | ssDNA | none | 1240K | - | 142 |
| S41 | *E. tonsillarum* | 1500±250 | Nepal | 91.53% | 4.2769× | 82.3213 | 3.99% | 3.94% | Human | Tooth | dsDNA | none | Shotgun | - | 169 |
| GCF004 | *E. rhusiopathiae* | 1425±75 | England | 2.53% | 0.0259× | 49.1061 | 22.81% | 20.10% | Human | Tooth | dsDNA | none | Shotgun | - | *Scheib et al (In preparation)* |
| GCF007 | *E. rhusiopathiae* | 1425±75 | England | 31.09% | 0.3865× | 44.3931 | 16.45% | 17.65% | Human | Tooth | dsDNA | none | Shotgun | - | *Scheib et al (In preparation)* |
| EDI001A | *E. rhusiopathiae* | 1400±50 | England | 1.47% | 0.0161× | 61.1049 | 2.25% | 5.26% | Human | Tooth | dsDNA | full | Capture | *Y. pestis* + HHV6 | <sup>6</sup> |
| EDI001 | *E. rhusiopathiae* | 1400±50 | England | 65.2% | 1.1931× | 43.571 | 13.87% | 13.80% | Human | Tooth | dsDNA | full | Shotgun | *Y. pestis* + HHV6 | *Scheib et al (In preparation)* |
| EDI012 | *E. rhusiopathiae* | 1400±50 | England | 87.99% | 4.5414× | 52.8114 | 29.97% | 29.87% | Human | Tooth | dsDNA | none | Shotgun | - | *Scheib et al (In preparation)* |
| EDI124 | *E. rhusiopathiae* | 1400±50 | England | 76.74% | 1.8945× | 44.4405 | 17.65% | 18.32% | Human | Tooth | dsDNA | none | Shotgun | - | *Scheib et al (In preparation)* |
| BMN020 | *E. rhusiopathiae* | 1400±20 | England | 5.66% | 0.0587× | 38.7338 | 25.04% | 20.43% | Human | Tooth | dsDNA | none | Shotgun | - | *Scheib et al (In preparation)* |
| ADN016 | *E. rhusiopathiae* | 1250±100 | Germany | 2.05% | 0.0187× | 47.2484 | 17.27% | 1.25% | Human | Petrous | ssDNA | none | 1240K | - | <sup>142</sup> |
| ADN010 | *E. rhusiopathiae* | 1250±100 | Germany | 2.92% | 0.0257× | 46.8014 | 14.89% | 0.73% | Human | Petrous | ssDNA | none | 1240K | - | <sup>142</sup> |
| ADN004 | *E. tonsillarum* | 1250±100 | Germany | 6.43% | 0.0681× | 48.9557 | 18.06% | 1.05% | Human | Tooth | ssDNA | none | 1240K | - | <sup>142</sup> |
| KIL026 | *E. rhusiopathiae* | 1000±350 | Ireland | 8.36% | 0.0882× | 52.5306 | 21.86% | 1.10% | Human | Tooth | ssDNA | none | Shotgun | - | <sup>142</sup> |
| KIL034 | *E. rhusiopathiae* | 1000±350 | Ireland | 91.92% | 30.058× | 60.964 | 11.83% | 1.95% | Human | Tooth | ssDNA | none | 1240K | - | <sup>142</sup> |
| KIL004 | *E. tonsillarum* | 1000±350 | Ireland | 17.83% | 0.2016× | 49.9257 | 21.63% | 1.32% | Human | Tooth | ssDNA | none | 1240K | - | <sup>142</sup> |
| V2T | *E. rhusiopathiae* | ~950 | UK | 1.23% | 0.0127× | 68.0881 | 14.47% | 5.88% | Human | Tooth | dsDNA | none | Shotgun | - | <sup>170</sup> |
| SWG009 | *E. tonsillarum* | 945±35 | Germany | 5.08% | 0.0528× | 57.1334 | 1.45% | 2.30% | Human | Tooth | ssDNA | none | 1240K | - | <sup>142</sup> |
| COM021 | *E. rhusiopathiae* | 800±200 | England | 11.15% | 0.1193× | 46.2285 | 13.99% | 13.60% | Human | Tooth | dsDNA | none | Shotgun | - | <sup>167</sup> |
| KPN009 | *E. rhusiopathiae* | 800±100 | Denmark | 2.52% | 0.0297× | 44.4076 | 25.46% | 0% | Human | Tooth | ssDNA | none | 1240K | - | <sup>142</sup> |
| A19x21 | *E. tonsillarum* | 622±227 | Denmark | 33.55% | 0.4426× | 71.875 | 10.06% | 11.52% | Human | Tooth | dsDNA | none | Capture | *Y. pestis* | <sup>119</sup> |
| ES8291 | *E. rhusiopathiae* | 599±1 | England | 2.05% | 0.0229× | 46.461 | 2.88% | 2.21% | Human | Tooth | dsDNA | full | Capture | *Y. pestis* | <sup>5</sup> |
| JDS004 | *E. rhusiopathiae* | 590±150 | England | 4.84% | 0.0629× | 61.5753 | 29.21% | 27.13% | Human | Tooth | dsDNA | none | Shotgun | - | <sup>167</sup> |
| STN020A | *E. rhusiopathiae* | 390±75 | Switzerland | 15.42% | 0.18991× | 56.282 | 1.32% | 0.81% | Human | Tooth | dsDNA | full | Capture | *Y. pestis* | <sup>8</sup> |
| STN004A | *E. rhusiopathiae* | 390±75 | Switzerland | 4.4% | 0.0446× | 57.916 | 0.26% | 0.88% | Human | Tooth | dsDNA | full | Capture | *Y. pestis* | <sup>8</sup> |
| JR135142 | *E. rhusiopathiae* | 335±8 | USA | 8.96% | 0.0961× | 104.95 | 11.49% | 0% | Dog | Tooth | dsDNA | full | Shotgun | - | <sup>123</sup> |
| JR68100 | *E. rhusiopathiae* | 335±8 | USA | 4.25% | 0.0439× | 78.186 | 0.8% | 20.78% | Dog | Tooth | dsDNA | full | Shotgun | - | <sup>123</sup> |
| JR75943 | *E. tonsillarum* | 335±8 | USA | 65.34% | 2.0968× | 108.69 | 1.09% | 5.5% | Dog | Tooth | dsDNA | full | Shotgun | - | <sup>123</sup> |
| F1364-1436 | *E. rhusiopathiae* | 298±1 | Spain | 71.69% | 1.5293× | 111.472 | 9.81% | 8.85% | Human | Tooth | dsDNA | none | Shotgun | - | <sup>106</sup> |
| F1691-1810 | *E. rhusiopathiae* | 298±1 | Spain | 26.81% | 0.3257× | 105.23 | 10.27% | 11.35% | Human | Tooth | dsDNA | none | Shotgun | *S. enterica* | <sup>106</sup> |
| HTC009 | *E. rhusiopathiae* | 100±10 | England | 93.95% | 15.013× | 55.1976 | 8.13% | 7.80% | Human | Tooth | dsDNA | none | Shotgun | - | <sup>167</sup> |
| Nr73 | *E. rhusiopathiae* | ~60 | Switzerland (*) | 2.16% | 0.0234× | 82.5308 | 2.83% | 0% | Human | Bone | ssDNA | none | Shotgun | - | <sup>168</sup> |
| Nr72 | *E. rhusiopathiae* | ~60 | Switzerland (*) | 3.7% | 0.0395× | 84.6816 | 0.62% | 0% | Human | Bone | ssDNA | none | Shotgun | - | <sup>168</sup> |
| AM71 | *E. rhusiopathiae* | ~60 | Switzerland (*) | 4.5% | 0.0488× | 86.6108 | 0.49% | 1.12% | Human | Bone | ssDNA | none | Shotgun | - | <sup>168</sup> |
| 15594A | *E. rhusiopathiae* | Missing | Missing | 90.99% | 10.26× | 70.1311 | 1.14% | 0.31% | Human | Missing | dsDNA | UDG | Shotgun | - | <sup>141</sup> |

\* Individuals originated in the Chilean Patagonia but died in Zurich.

We validated the presence of 60 previously unreported *Erysipelothrix spp.* genomes in published data (45 *E. rhusiopathiae* and 15 *E. tonsillarum*), with depths of coverage ranging from 0.01× to 65.38× (Average of 3.998×, Median of 0.15×) [Table 2]. Damage patterns match expectations for authentic ancient DNA in regard to library type (single stranded - ss versus double stranded - ds) and enzymatic treatment (UDG versus non-UDG), with terminal C>T deamination damage ranging between 0.62–45.46% for non-UDG dsDNA library data. *Erysipelothrix spp.* genomes have been retrieved from at least 4 different sources including teeth, petrous bone, post-cranial elements and sediment. Due to the fact *Erysipelothrix tonsillarum* has been found in sediments associated with human and animal inhabitation (EMN005)^130^, we decided to consider the species as potentially recovered from the environment. Contrary to this, we found no traces of *E. rhusiopathiae* in archaeological sediments. We find *Erysipelothrix spp.* sequences in data from different library construction approaches such as shotgun libraries and *Y. pestis, S. enterica,* and 1240k captures. From a 1240k capture originating from an Irish human individual dating back to the 10th century (KIL034), we reconstructed a high quality genome at a depth of 30.058×^142^.

For *Erysipelothrix* associated with *Y. pestis* capture, we analysed 5 additional shotgun libraries from different skeletal elements of EDI001, an Anglo-Saxon individual for which we detected *E. rhusiopathiae* in published *Y. pestis* capture data. We detected more than 1,000 sequences mapped against the bacterium in EDI001 in both capture and screening data originating from tooth root, but none in libraries generated from maxilla or dentin powder [see Supplementary Information SI5 and SI6; Supplementary Figure S14]. We additionally analysed which genomic features of *E. rhusiopathiae* display affinity to *Y. pestis* and *S. enterica* sequences. *E. rhusiopatiae* genomes recovered from pathogen capture present higher read density in 4 regions associated with conserved genes (r/tRNA), but display a similar coverage distribution to shotgun samples otherwise [Supplementary Figure S15]. We have discarded those regions in downstream analysis and considered the genomes from capture data as suitable for phylogenetic projection.

For the 184 previously published *Y. pestis* samples we compared the presence of *Erysipelothrix spp.* at different points in time (Supplementary Table S7). We found that *Erysipelothrix spp.* were present in 4 of 93 samples (4.3%) in the Second Pandemic (SP) samples and 1 of 35 samples (2.9%) in the First Pandemic samples (FP), whereas prevalence was markedly higher in Prehistoric (Pr) samples (10/46; 21.7%). Pairwise Fisher’s exact tests showed that prevalence in Pr was significantly higher than in SP (*p* = 0.0024) and FP (*p* = 0.0196), while no difference was observed between SP and FP (*p* = 1.00). These results could be affected by the overwhelming proportion of captured genomes of historical *Y. pestis* (in contrast to prehistoric genomes that originate from shotgun libraries in the majority of cases), hindering our ability to detect *Erysipelothrix spp.* bacteria. Within individual periods, no significant differences in prevalence were detected between capture and shotgun libraries (SP: 3/73 vs 1/20, Fisher’s exact test *p* = 1.00; FP: 1/32 vs 0/3, *p* = 1.00; Pr: 4/23 vs 6/23, *p* = 0.72). Some caution should be exercised in regard to these results, given the low sample size.

### Ancient Erysipelothrix rhusiopathiae genetic affinities

*E. rhusiopathiae* is characterised by the high level of recombination present in its phylogeny (up to 56% of the core genome)^50^. We explored recombination between ancient and modern *E. rhusiopathiae* with ClonalFrameML^143^, identifying 9,369 of such events across 196 modern and 8 high coverage (>4×) ancient genomes, accounting for a total of 1,241,692 bp (approximately 70% of the genome) [Supplementary Figure S15; Supplementary Table S8].

We built a phylogenetic SNPs dataset, excluding repetitive regions, conserved genes (t/rRNAs), low quality SNPs, and recombinant tracks that could potentially affect the tree topology [See Methods][Supplementary Table S8]. This high quality dataset included 6,809 SNPs identified in the previous 170 modern and ancient *E. rhusiopathiae* genomes. Those high quality SNPs were called on 12 *E. tonsillarum* strains, 3 *Es2,* and 5 ancient moderate-coverage genomes (between 4× and 1×). We built a ML phylogeny using raxml-ng and this high quality dataset, with 1,000 bootstraps and the GTR+G model. We used epa-ng to project low coverage genomes (between 1× and 0.05×) onto the tree, with an average first placing Likelihood Weight Ratio (fpLWR) of 80.96%, only displaying those genomes with a fpLWR above 70% [Figure 4; Supplementary Table S8; Supplementary Figure S17]. The tree displays clear differentiation of the 4 main clades (average TBE=0.7124; average FBP=52.6955). Most of the ancient samples cluster at the base of the tree in Clade 2, with the older samples occupying deeper positions in the phylogeny, but without a clear pattern regarding geographical order nor host. Exceptions to this behaviour are TGEZ06 that is projected at the base of *E. rhusiopathiae* diversity (fpLWR=94.97%), the early modern genomes HTC009 and F1364-1436 positioned at the base of Clade 3 (TBE of 0.9998 and 0.9901 respectively), and the rest of the low coverage early modern genomes (F1691-1810 and JR135142) projected at either the base or within Clade 3. GSP013 is robustly positioned among other Neolithic and Bronze Age samples within Clade 2 (fpLWR= 99.72%).

**Figure 4.**
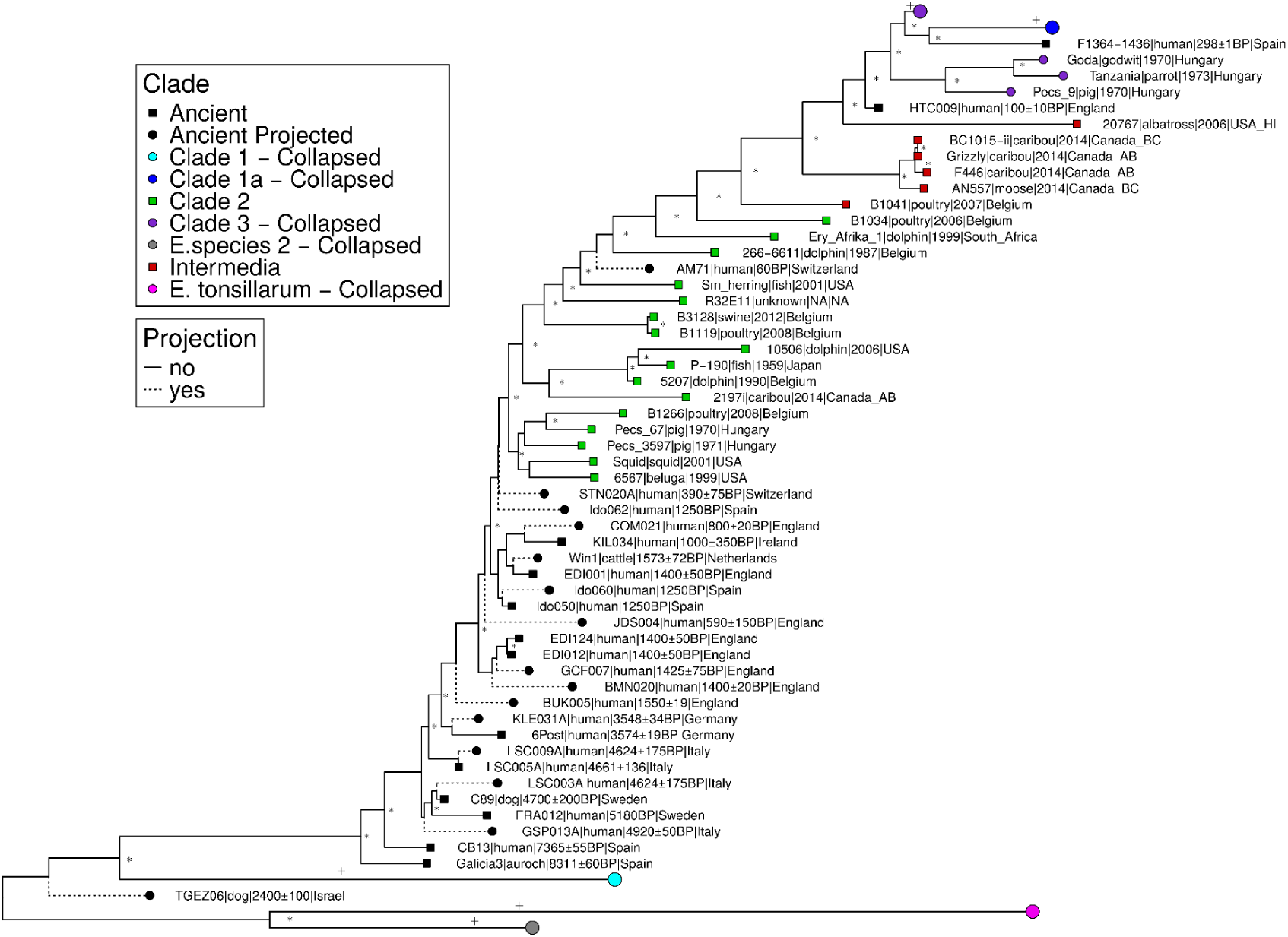
Phylogeny of modern and ancient *Eryspelothrix rhusiopathiae*. ML tree of 6,809 recombination free SNPs found in 224 genomes of modern and ancient *E. rhusiopathiae.* Low coverage samples projected with epa-ng and with a first placement LWR≥70% are displayed with a dotted line. Nodes with a TBE≥0.8 are marked with a **\***. Collapsed nodes with a TBE≥0.8 are marked with a +.

In parallel, we created a high quality tree of *E. tonsillarum* using the same parameters, and we included C88, *E. rhusiopathiae* and *E. sp. Strain 2*. A ML tree was built with raxml-ng (average TBE=0.701; average FBP=65.933), and low coverage genomes were again projected using epa-ng (average fpLWR=90.83%). All ancient genomes recovered cluster within the modern diversity of *E. tonsillarum* [Supplementary Figure S18].

To identify variation in gene content between ancient (13 genomes with coverage above 1× and GSP013) and modern genomes of *E. rhusiopathiae* that could explain changes in pathogenicity, we examined the coverage of a set of 45 previously described virulence genes^133,144^ [Figure 5A, Supplementary Table S9]. Ancient genomes displayed patterns similar to those of Clade 1, Clade 2 and Intermedia genomes, their closest phylogenetic relatives. Among the examined virulence-associated loci, ERH_1467, which encodes a biofilm-associated surface protein^145,146^, exhibited variable coverage across all samples [Figure 5A]. Most genomes retained high read depth for this locus, whereas Medieval genomes, FRA012 and Galicia3 showed a reduced or absent signal, suggesting possible gene loss. The lysophospholipase C gene (ERH_1433), previously identified in the Fujisawa genome as part of a lysophospholipase A/B/C cluster^147^, was also absent in 2 Medieval genomes (EDI012 and EDI124), 6Post and GSP013. Lysophospholipases can modulate host-cell membrane composition and signaling, thereby influencing intracellular survival and immune responses^148^. ERH_0278, predicted to encode a surface protein of unknown function^145^, was likewise undetected or non-covered in all Neolithic-Bronze Age genomes (including GSP013) and absent or truncated in all Medieval samples. Given its predicted surface localization, it may act as an adhesin or immune-interactive factor. ERH_1258 (uncharacterized surface protein) was absent in Galicia3, Medieval genomes and in several Clade 1 genomes. ERH_1472 is an internalin-like surface protein ^145^. These proteins are known to mediate host-cell attachment and invasion. ERH_1472 was absent in FRA012 and Medieval genomes, in addition to most Clade 3 strains. The collagen-binding protein ERH_1436^145^ was broadly conserved but variably covered among human isolates, with absence in Medieval genomes and 6Post. Such heterogeneity is consistent with prior findings that *E. rhusiopathiae* collagen-binding adhesins diversify across lineages, a pattern likely driven by host or tissue-specific pressures ^144,149^.

**Figure 5.**
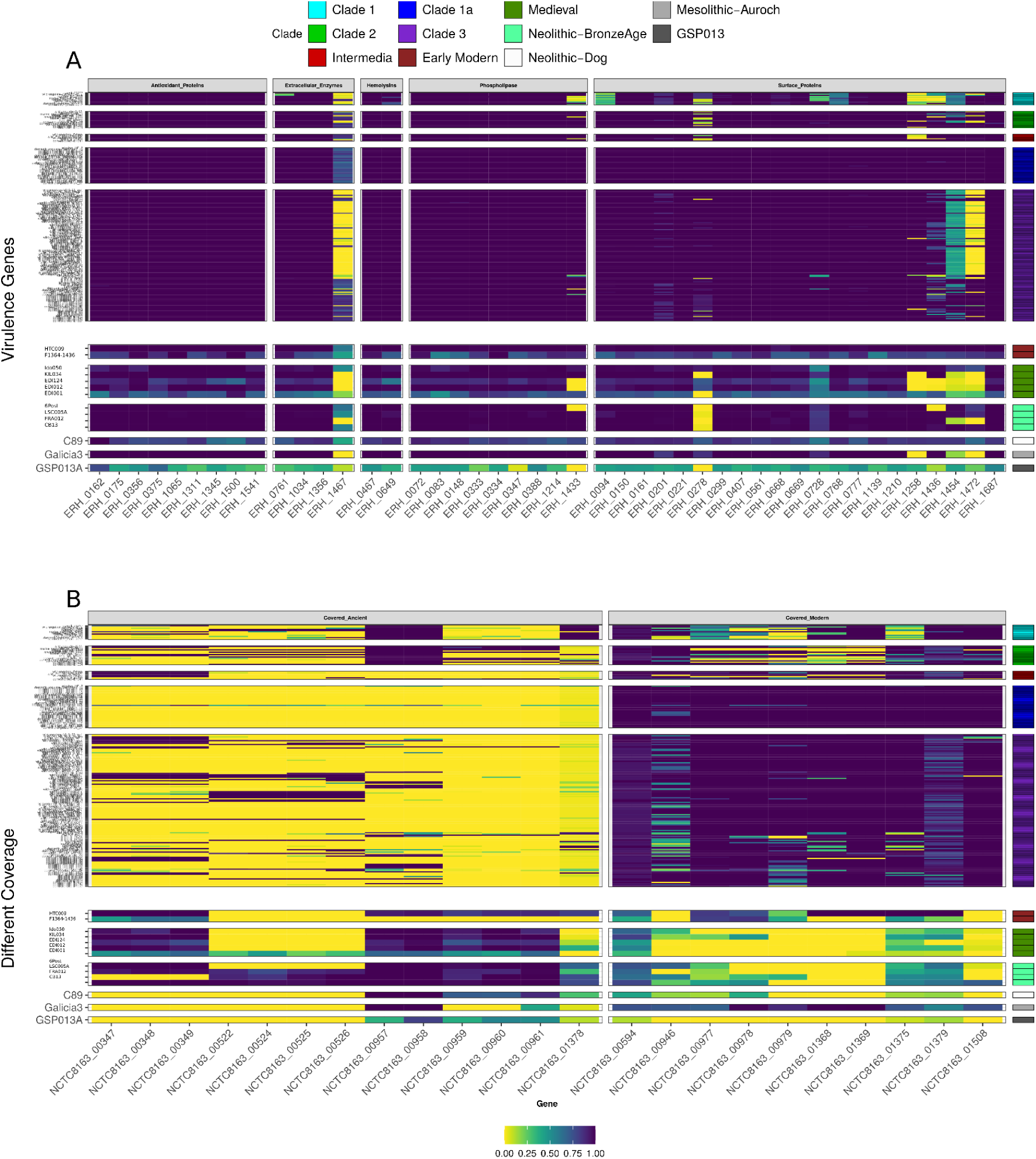
Gene absence/presence between ancient and modern *E. rhusiopathiae* genomes. The upper panel focuses on virulence genes described in the literature. The lower panel describe genes screened in this study (Absent in modern and present in ancient; present in modern and absent in ancient; mostly present in modern and absent in ancient, and truncated in ancient). Colour bars on the right site indicate the Clade.

In addition to known virulence genes, we explored the coverage of 23 genes found in regions which displayed different coverage patterns between ancient and modern genomes [Figure 5B, Supplementary Table S9]. Ancient genomes and specifically those retrieved from humans present three genomic regions largely absent from most present day genomes. The first one is a ∼3kb region at position 347,330 of *E. rhusiopathiae* chromosome (present in 20% of the modern and in 72% of the ancient human genomes), comprising three genes (loci NCTC8163_00347-49) encoding a colanic biosynthesis UDP-glucose lipid carrier transferase (WcaJ), a diaminopimelate decarboxylase (DAPDC), and a UDP-glucose 4, 6-dehydratase (RmlB). All the mentioned genes are involved in synthesis of the peptidoglycan cell wall, as well as exopolysaccharides and lipopolysaccharides^150–154^. The second is a 5kb region at position 541,590, containing 5 genes (NCTC8163_00522-26) missing in most modern strains and present in the 3 Neolithic human genomes. This region contains a GNAT acetyltransferase(00522), a MazG-like family (00524), EcoKMrr (00525), and 2 Uncharacterised proteins. GNAT acetyltransferase activity is linked to virulence in different bacteria genii^155^, and transcriptional and protein synthesis regulation in bacteria under me^156^. MazG proteins have been shown to mediate cellular death and promote survival under nutritional derived stress ^157^. Mrr is a restriction gene that attacks foreign methylated DNA and has been linked to bacteriophage protection and response to environmental stress in *E. coli*^158^. The third region mostly covered in ancient human genomes but not in modern ones, is a 6kb segment at position 976,063, containing genes encoding for a HTH domain protein(CTC8163_00957), a IS3 family transposase (CTC8163_00958), the Internalin-J precursor (InlJ) (gene NCTC8163_00959), an Endonuclease-Exonuclease-phosphatase domain superfamily protein (EEP-CTC8163_00961) and an uncharacterised protein (NCTC8163_0095960; present in 6.1% of modern, 91.1% of ancient human genomes). InlJ functions as an adhesin and is a known virulence factor in *Listeria monocygotenes,* where it facilitates adhesion to mammalian cells^159^, and contributes to biofilm formation in *Porphyromonas gingivalis*^160^. Proteins from the EEP family have been linked to host immune system evasion through degradation of secreted extracellular traps^161,162^.

In regard to genes absent in ancient genomes, all ancient human genomes and C89 lack loci NCTC8163_01369 (macB; Macrolide export ABC transporters), Notably, one ancient wild ungulate genome (Galicia3) exhibits clear coverage across macB, suggesting that this efflux module was not universally absent in the pre-antibiotic era. This pattern likely reflects host-or lineage-specific retention or acquisition rather than a simple modern-only gain driven by selective pressure exerted by antibiotics. Other patterns for antibiotic resistance have been observed in a region comprising 3 genes (present in 30% of modern versus 0% of ancient) NCTC8163_01354 (ABC transporter permease subunit), NCTC8163_01355 (DrrA; Daunorubicin/doxorubicin resistance ATP-binding protein), NCTC8163_01356 (YkoD; Putative HMP/thiamine import ATP-binding protein), which are linked to resistance to different antibiotics^163–166^ [Supplementary Table S11].

Given the observed pattern, we analysed population stratification among the different *E. rhusiopathiae* Clades with both a wide genome and virulence gene SNP PCA [Supplementary Figure S19]. The results show that Prehistoric and Medieval genomes cluster with Clade 2 in both cases, while 17th and 19th century genomes (1364-1436 and HTC009) cluster with Intermedia and Clade 3 strains. We run genome wide Fst scans between Ancient genomes (Prehistoric and Medieval) and Clade 2 strains, detecting 7 regions with high Fst [Supplementary Figure S20]. The top 1% Fst genes in those regions contain genes associated with amino-acid and lipid metabolisms, and wall synthesis [Supplementary Table S10].

### Discussion

We have reported newly generated genome-wide data for 8 Copper Age individuals from the Italian site of *Grotta della Spinosa*. The genetic ancestry of these individuals aligns with the diversity of Neolithic populations from the Italian peninsula, suggesting that they were local. Notably, we found that one individual (GSP013), an adolescent female of around 16–18 years of age at death, had both aDNA sequences belonging to *Y. pestis* (for which we retrieved a complete genome), *HBV,* and to the generalist zoonotic bacterium *E. rhusiopathiae*, suggesting a possible case of co-infection. In addition to that, 3 other of the 8 individuals from *Grotta della Spinosa* show molecular evidence for the presence of *HBV*, suggesting an endemic burden for this disease. Finally, we found evidence of bacteria species from the genus *Erysipelothrix* in at least 60 published ancient individuals, including 51 humans^5,6,8,13,16,23,24,84,88,89,92,105,106,119,126,128,141,142,167–170^, 6 dogs^122,123^, and 3 bovids^125,126^.

### Yersinia pestis in Southern Europe

Our data confirms a case of *Y. pestis* infection in Western Europe dated 300 to 400 years before other geographically close genomes from Central Europe (VLI092 and GRS004)^23^. Phylogenetically, the strain falls within the LNBA-clade basal to most previously known European genomes, and only slightly more derived than the ones from the Northern Caucasus and the Altai region. Virulence gene screening confirms that the strain in GSP013 lacks three key genes, *ymt*, YPMT1.66c and the filamentous prophage. The *ymt* gene is essential for flea-borne transmission^94^ as it protects *Y. pestis* against toxic by-products of blood digestion and contributes to biofilm stability, thereby enabling proventricular blockage in rodent fleas and efficient transmission to mammals^171,172^. However, according to the Unified Theory of Flea-Borne Transmission^94^, *Y. pestis* genomes lacking *ymt* can still be transmitted by fleas via early-phase transmission, although its efficiency remains debated in the literature^173–176^. Previous works have shown that *ymt*(–) *Y. pestis* can be transmitted by human ectoparasites, such as body lice^177^, suggesting that transcutaneous transmission of plague could occur even in the absence of the *ymt* gene. This implies that bubonic forms of the disease might have already been present in prehistoric times. However, the absence of YPMT1.66c would reduce the ability of *Y. pestis* to survive within macrophages and replicate efficiently at the flea bite site, which are essential steps for the establishment of bubonic plague^97^. Consequently, the emergence of fully developed bubonic forms during the LNBA period remains unlikely, despite the possibility of transmission via ectoparasites^23^. The loss of the filamentous prophage has been associated with reduced bacterial virulence and fitness during mammalian infection, raising questions of the pathogenic potential of prehistoric *Y. pestis* genomes^22,93,178,179^. Another major virulence determinant, the protease *pla*, which is located on the pPCP1 plasmid in the vast majority of modern strains^99,180^, also provides insights into pathogenicity. In ancestral lineages, *pla* carries a single isoleucine substitution at position 259 (I259) located on the surface loop 5, whereas modern lineages have a conserved threonine at this residue (T259). Experimental work suggests that genomes with the ancestral variant are less efficient in colonizing tissues, but can cause the pneumonic form of plague^88,181^. This distinction might suggest that early *Y. pestis* genomes, such as the one carried by GSP013, could have also caused deadly contagious respiratory infections. When considered together with the lack of evidence for plague-related mass mortality at *Grotta della Spinosa*, these findings suggest that non-bubonic manifestation of plague may have predominated. Such forms are generally less likely to cause a major epidemic than the bubonic one, but the epidemiological impact still warrants further investigation. This early case of a *Y. pestis* infection in southwestern Europe, far from the predicted historical focus of the disease in the Eurasian steppe, in conjunction with its gene content, is in agreement with the hypothesis that *Y. pestis* diversity was already established in the continent, mainly found in animal reservoirs^23^. This also hints at a possible zoonotic transmission of Late Neolithic–Bronze Age plague^14^.

### Zoonotic context and implications

The presence of the zoonotic pathogens *E. rhusiopathiae* and *E. tonsillarum* in prehistoric individuals coinfected with *Y. pestis* strengthens the hypothesis that prehistoric genomes of *Y. pestis* could have been transmitted by contact with animal intermediate hosts or their waste^16,22,34^. This is also reflected by the coexistence in time and space of *Erysipelothrix* at archaeological sites with human individuals infected by other potential zoonotic pathogens, such as *Frälsegården* (FRA - *Yersinia enterocolitica* and *Borrelia spp.)*, *Hagios Charalambos* (HGC - *Salmonella enterica*), *Murzihinskiy* (MUR - *Salmonella enterica*), *la Sagrera* (F1691-1810 - *Salmonella enterica*). Alternatively, the presence of *Y. pestis* could have created a ‘permissive environment’ for other, chronic pathogens as discussed in Guellil et al. 2022^121^.

Despite the common misconception, experimental data show that *E. rhusiopathiae* does not grow in soil, but soil can be contaminated through infected animals^38^. *E. rhusiopathiae* can, however, survive in soil for up to 35 days^48^, comparable to other zoonotic bacteria often analysed in aDNA studies, such as *Yersinia pestis* (24 days)^182^, or *Salmonella enterica* (216 days)^183^. This seems to agree with the results of archaeological sediment screening, in which it appears unlikely that *E. rhusiopathiae* is a common, widespread soil contaminant. In contrast, *E. tonsillarum* has been identified in human associated sediments^130^, reflecting its potential as an environmental contaminant, or at least, its potential to be recovered in sediments from environments in which humans or animals were present. Nonetheless, deamination damage patterns are in line with the authenticity of those sequences, suggesting they are at least from an ancient origin. We also present suggestive evidence of a septicemic manifestation of the bacteria in EDI001 and GSP013 [see Supplementary Information SI6]. More studies should focus on this aspect in the future to properly validate this hypothesis.

Overall, both the different sequence characteristics and inferred phylogenetic placing point to the fact that the retrieved genomes are indeed ancient, although whether the bacteria infected the analysed individuals during life, or only contaminated them later on, is difficult to determine^127^. The disease can produce high mortality if left untreated, especially in the pre-antibiotic era (with a mortality of 38% with antibiotic treatment versus 100% without), mainly through endocarditis complications^49,184^. Sampling biases may also have affected our analyses, since most of the sequencing data of the Medieval individuals included here were generated using *Y. pestis* capture^5,6,8,119^, hindering our ability to properly detect the presence of *Erysipelothrix.* We nevertheless show that sequences originating from capture and shotgun libraries from the same individual are comparable (EDI001). Yet, most individuals with shotgun and capture libraries present *Erysipelothrix* sequences only in the shotgun libraries, indicating the possibility of a substantial proportion of undetected cases in First and Second Plague Pandemics samples with only capture data. Nonetheless, we speculate that the overrepresentation of *Y. pestis* and *Erysipelothrix* co-infections during prehistory (compared to historical times) could be associated with the Neolithic transition, which increased close contact between human and domestic animals due to the introduction of husbandry, sedentism and agriculture, facilitating the transmission of these bacteria between both^22,105^.

### 7,000 years of Erysipelothrix infections in humans

*Ersypelothrix spp.* have infected animals for at least 1 million years^185^, and, until our findings, the only cases reported for ancient humans was an individual from Russia 4,000 years ago^37^, and 3 Iberian individuals dating back to the 10th century of the common era ^36^. Our analysis found 45 new cases of *E. rhusiopathiae,* pushing back the first evidence for a possible human infection to a Neolithic human individual from Spain (CB13 - 7,365 years calBP)^128^. We report seven more high coverage ancient genomes of *E. rhusiopathiae* retrieved from an aurochs (8,311 years BP)^125^, a Neolithic individual from Scandinavia (FRA012 - 5,200 BP)^24^, a Copper Age individual from *La Sassa* in Italy (LSC 4,661±136 calBP)^84^, a Bronze Age human individual from Germany (6Post, 3,574 years BP)^23^, a Medieval individual from Ireland (KIL034 1,000±350BP)^142^, an English individual from the 19th century (HTC 100±10BP), and a British Individual without metadata (15594A)^141^. The phylogenetic analysis of *E. rhusiopathiae* and *E. tonsillarum* supports their role as generalist pathogens in prehistoric times, since they do not fall into clades with known host specificity. Although projected phylogenetic placement of low coverage genomes have to be taken with caution, prehistoric genomes of *E. rhusiopathiae* fall basal to modern Clade 2, without evidence of clonal expansion characteristic of epizootic outbreaks in Arctic muskox^40,144^, porpoises in the Netherlands, or swine in Japan^186^ of Clade 3 and Clade 1a. The former tree structure of Clade 2 and ancient genomes suggests frequent infection. Genomically speaking, the ancient *E. rhusiopathiae* genomes are similar to extant genomes of the bacterium. Specifically, all analysed ancient genomes seem to have comparable gene content, and phylogenetic affinities, to current day strains from Clade 2 and Intermedia. An exception to this is the presence of genes WcaJ, DAPDC, RmlB (involved in the synthesis of the peptidoglycan wall)^150–154^, InlJ (cell adhesion and biofilm formation)^160,187^, and EEP (immune system evasion)^162^, absent in most modern genomes, and that seem to point to differences of infectivity between ancient and modern genomes. Conversely, ancient genomes do not present coverage of genes macB, DrrA, YkoD, and an unnamed ABC transporter permease (NCTC8163_01354), all of them linked to antibiotic drug resistance^163–166^, that could be explained by the lack of selective pressure created by the use of antibiotics^188^, which has been observed in other ancient pathogens^189,190^.

These new results provide a broader insight into the spatiotemporal boundaries of *Y. pestis* in prehistoric Europe. Additionally, they highlight the historical importance of zoonotic pathogens such as *E. rhusiopathiae*, whose role in human diseases has been historically neglected. More studies involving ancient and modern genomes of these bacteria are necessary to have a better understanding of its adaptation and diversification as a pathogen.

## Methods

### Archaeological context of the Grotta della Spinosa, Italy

*Grotta della Spinosa* (Grosseto, Tuscany, Italy) is a natural cavity in the travertine bank near Torrente Gavosa in Massa Marittima. The site was excavated between 2001–2003 by the Superintendency for Archaeological Heritage of Tuscany under the supervision of Biancamaria Aranguren and Elsa Pacciani.

The cavity consists of a single chamber, approximately 12 x 6 m in maximum dimensions. The archaeological deposits had been disturbed by looters across approximately 60 % of the cave, however, the remaining intact portions amount to approximately 80 square meters and allow reconstruction of the stratigraphy of the cave’s use. Above the sterile base soil there was a layer (US 9), preserved across the entire walkable surface of the cave, without any burials, with traces of hearths, animal bones, lithic and bone tools and ceramic fragments which testify to the residential use of the cave in the second half of the 5th millennium BC, during the Middle Neolithic (attributed to the Middle Tyrrhenian facies with contacts with Serra d’Alto-Diana and Chassey cultures)^192,193^. The above layer (US 8) contained a dense deposit of disarticulated and fragmented human remains along with artefacts such as stone and bone tools, ornamental objects, pottery fragments, and copper daggers. Typological assessments of the material culture place these burials in the Middle Copper Age, consistent with the cultural phase of Sassi Neri^53^, alongside two Remedello-type daggers^53^. These depositions were sealed by a layer (US 7) composed of debris. *Grotta della Spinosa* therefore contains a partially preserved Copper Age funerary deposit, where the position of the human remains and related material culture could be considered the result of intentional interventions resulting from complex funerary and post-depositional manipulations of the remains. [Supplementary Figure S2]. For this study, we collected seven samples from human remains from different areas of the same stratigraphic unit (US 8), plus two more samples from layers 4 and 5 [Supplemental Table S1A].

### DNA extraction and sequencing

The aDNA workflow was performed at the state-of-the-art ancient DNA laboratory at the Institute of Genomics, University of Tartu (Estonia). For each sample except for sample GSP005, a portion of the skeletal human remains was sampled and one extraction as well as one dsDNA library were built. For GSP005, two extracts and two dsDNA libraries were produced, one from the petrous bone and one from the incus.

Portions of the crown of the teeth, tooth roots, petrous cores, and an incus were sampled with a sterile drill wheel and briefly brushed to remove surface dirt with full strength household bleach (6% w/v NaOCl) using a disposable toothbrush that was soaked in 6% (w/v) bleach prior to use. Contaminants were removed from the surface of the sampled area by soaking in 6% bleach for 5 minutes, then rinsing three times with milli-Q water (Millipore) and lastly soaking in 70% ethanol for 2 minutes, shaking the tubes during each round to dislodge particles. Lastly, the sample was left to dry under UV light for 2 hours. Next, a buffer of 2ml/100 mg sample weight of 0.5M EDTA Buffer pH 8.0 (Fluka) and 50 μl/100 mg sample weight of Proteinase K 10 mg/ml (Roche) was added and the sample was left to digest for 72 hours at room temperature on a nutating mixer. ^194,195^

The DNA solution was concentrated to 250 μl using Vivaspin Turbo 15, 30,000 MWCO PES, Sartorius concentrators and purified in a large volume column (High Pure Viral Nucleic Acid Large Volume Kit, Roche) using 2.5 ml of PB buffer, 1 ml of PE buffer and 100 μl of EB buffer (MinElute PCR Purification Kit, QIAGEN).

Only one extraction was performed per sample and 30μl were used to prepare dual-index double-stranded libraries. The dsDNA libraries were built following established protocols ^196,197^. Three verification steps were implemented to make sure library preparation was successful and to measure the concentration of DNA in the libraries - fluorometric quantitation (Qubit, Thermo Fisher Scientific), parallel capillary electrophoresis (Tape Station, Agilent Technologies) and qPCR (KAPA Library Quantification Kit - Illumina^®^ platforms).

DNA was initially shotgun-sequenced using the Illumina NextSeq500/550 High-Output paired-end 150 cycle kit at the University of Tartu Institute of Genomics Core Facility. The deeper sequencing (GSP002, GSP004 and non-enriched library of GSP013) was performed in the same facility using the Illumina NextSeq2000 sequencing platform (P4 100 Kit). Following amplification, the enriched library of GSP013 was sequenced on the same Illumina NextSeq2000 sequencing platform (P2 100 Kit), with other samples.

### Target enrichment

The dsDNA library of GSP013 was enriched using a custom *Yersinia pestis/Yersinia pseudotuberculosis* myBaits target enrichment kit from Daicel Arbor Biosciences (v5)^10^ . The design comprises the *Y. pestis* CO92 reference sequence (including all plasmids) and the *Y. pseudotuberculosis* reference sequence (NC_006155.1). In order to reach the desired concentration, the library was re-amplified using KAPA Hifi Hotstart ReadyMix (2X) and primers IS5 and IS6 51 prior to enrichment ^198^ . The sample was enriched in a single reaction using 5 µl of baits. Hybridization was carried out for 35 h at 62 °C. The enriched library was amplified using KAPA HiFi HotStart ReadyMix (2X) DNA Polymerase and primers IS5 and IS6 51.

### Human sequence mapping and quality control

Due to the specifics of the NextSeq 500 technology, first, the sequences of the adapters, indexes, and poly-G tails were removed using cutadapt-1.11^199^. To avoid random mapping of sequences belonging to other species, sequences shorter than 30 bp were filtered using the same program. The trimmed sequences were aligned to the human reference genome GRCh37 (hs37d5) using Burrows-Wheeler Aligner (BWA v0.7.12)^200^ using the command *mem* with re-seeding disabled. After the alignment, the human sequences were converted to the BAM format using samtools 1.3^201^. In case of paired-end sequencing, the different flow cell lanes were merged and duplicates were removed using MarkDuplicates by picard v1.2 (https://broadinstitute.github.io/picard/index.html) and samtools 1.3. Finally, indels were re-aligned using GATK 3.5^202^ and reads with a mapping quality less than 10 were filtered out using samtools v1.3. After mapping each sequencing run separately, the different runs and extractions of one sample were merged using MarkDuplicates by Picard v2.20 and samtools v1.9.

### aDNA authentication and sequencing statistics

To validate the presence of ancient DNA reads, we used mapDamage v2.0 to estimate the frequency of C => T substitutions at the 5′ ends of sequences due to cytosine deamination and the distribution of short reads^203,204^. Rates of modern human contamination for the mtDNA were estimated using ContamMix v1.0.11^205^. For male individuals, the contamination was estimated on the X chromosome using ANGSD-0.917^57^ command–doHaploCall [Supplementary Table S1C]. The samtools option *stats* was used to determine the number of (final) reads and average read length. The unique endogenous human DNA content was estimated based on the final reads (MQ10) divided by the number of trimmed reads. The genetic sex of each sample was estimated following a method described in^52^. The mtDNA haplogroup of each sample was estimated using haplogrep2 command line^206,207^ [Supplementary Table S1C]. The Y chromosome haplotypes were estimated using an in-house script.

### Identifying genetically identical individuals

To estimate the minimum number of individuals sampled from this disarticulated site for this study, we first estimated the possible genetic relationships using READv2^52,208^ between all the sequenced dsDNA libraries keeping in mind that a small samples set shifts the P0 towards a false negative. After that, we subsetted the sample set to samples above 0.03X of average human coverage, then used READv2 and the uniparental markers to confirm the minimum number of 8 unique individuals. The dsDNA libraries of the samples GSP005 and GSP013 were merged, as the samples were identified to stem from the same individual.

### Preparation of the human population genetics dataset

After merging the different sequencing runs to obtain higher average coverages and merge of identical individuals, we called the autosomal variants using ANGSD-0.916 ^57^ command *–doHaploCal*l using all the ‘1240k’ positions extracted from the binary genotype file (.bim) of the Allen Ancient DNA Resource (AADR) dataset. The ANGSD output (haplo.gz) was converted to plink format using PLINK v1.9 ^209^ and the autosomal SNPs were estimated using the command *–missing*. We selected six individuals with more than 20,000 autosomal ‘1240k’ SNPs for genome-wide analysis. We merged our newly generated dataset with the published available AADR dataset (v62.0, https://dataverse.harvard.edu/dataset.xhtml?persistentId=doi:10.7910/DVN/FFIDCW, version: January 2025) ^86^ subsetted to the HO array to increase the number of published present-day Eurasia populations keeping positions that are present at MAF > 0.1 in the dataset.

### Principal Component Analysis

We first converted the plink files to EIGENSTRAT format using convertf from the EIGENSOFT 7.2.0 package with the parameter familyname:NO and used the program smartpca with the parameter autoshrink:YES to project the newly generated and published ancient individuals onto the genetic variations of present-day Western Eurasian populations. The PC1 and PC2 were plotted using R.

### Metagenomic screening and sequence authentication

For metagenomic analyses, raw sequences were pre-processed using AdapterRemoval v2^210^, trimming adapters, and collapsing paired-endpair ended reads when necessary. Additionally, we discarded sequences under 30 bp in length, trimmed bases with qualities under 20, and trimmed Ns. Following this initial pre-prossencing, we used prinseq-lite to remove sequences with a complexity dust value under 7^211^, and then, deduplicated sequences using BBmap dedupe.sh^212^.

High-complexity deduplicated sequences were screened using KrakenUniq against the standard MicrobialDB (16/08/2020) ^213^. We filtered the resultant reports using an E-score given by the *kmers* per sequence divided by the genome coverage ^121^, with a minimum score of 7, and a minimum number of assigned reads above 100 (except for viruses).

### Yersinia pestis phylogenetic analysis

Ancient published *Yersinia pestis* genomes were mapped against *Y. pestis* CO92 reference genome using BWA backtrack with the same parameters as described above (-l 16500 -n 0.01 -o 2)^200,214^. Read duplicates were removed with picard MarkDuplicates^215^ and mapping quality 30 reads were keeped with samtool^203^. We used MapDamage2 to calibrate qualities and remap de recalibrated fastqs^10,14^. Following this we used *GATK Unified Genotyper*^202^ (*--emit-all-sites)* to generate callings of the mapped genomes. In order to create a high quality biallelic positions dataset, we filtered all calls using vcftools (including GSP013)^216^ for low quality positions, with a minimum depth of coverage of 3×, a minimum genotyping quality of 30, removing indels and heterozygous positions, maximum missingness per site of 40%, allele depth ratio of 90%, and a minimum physical distance of 3bp between SNPs. We also discarded t/rRNA regions and repetitive regions discovered with Dustmasker (>30bp)^217^. Variant positions found in the dataset were called from the *Y. pseudotuberculosis* genome to avoid long branches. This dataset contained a total of 2,026 SNPs across the 36 ancient genomes of *Y. pestis, Y. pestis* strain CO92, and *Y. pseudotuberculosis*

Additionally, to test bias introduced by terminal aDNA damage and errors we created 2 additional datasets. In one we removed singletos, while in the other we removed all C ↔ T and G ↔ A transitions. Those accounted for 961 and 674 SNPs respectively.

We used raxml-ng with the GTR+G model with 1,000 bootstraps to create a Maximum Likelihood phylogeny^91^. Support of the tree was assessed using Felsenstein’s bootstrap proportions (FBP) and Transfer Bootstrap Expectation (TBE). The final tree was visualised using R package ggtree^218^. Additional trees were visualised with iTOL^219^.

### Erysipelothrix rhusiopathiae phylogenetic analysis

All *E. rhusiopathiae* genomes were processed with AdapterRemoval^210^ with the aftermention parameters Modern *E. rhusiopathiae* genomes were mapped against *E. rhusiopathiae* reference genome NCTC8163 using *bwa mem* with default parameters^220^, while ancient genomes were mapped using *bwa backtrack* with parameters adapted for aDNA samples (-l 16500 -n 0.01 -o 2)^200,214^. Duplicates were removed with *picard MarkDuplicates*^215^, and reads with a mapping quality below 30 were filtered out^221^.

We generated VCF files using GATK *Unified Genotyper* with the option “*--emit-all-sites”* for each of the 196 modern *E. rhusiopathiae* and the 8 ancient high coverage ones (FRA012, HTC009, Galicia3, CB13, EDI012, HTC009, KIL034, LSC005A) mapped against the *E. rhusiopathiae* reference genome NCTC8163NCTC8163. We filtered them using vcftools for positions under 5 folds of coverage, genotyping quality under 20, and indels *(--minDP 5 --minGQ 20 --remove-indels).* With an inhouse script, we also reassigned the genotype of heterozygous positions if the majoritary allele was supported by 90% of the reads or more (Allele Depth filter), discarding non-reclassifiable positions. All the discarded positions per sample in those steps were appended and converted into sample specific bed files. Additionally, we screened the Erysipelothrix *rhusiopathiae strain NCTC8163* reference genome for the presence of repetitive regions (>30bp) using Dustmasker. We have also selected tRNA and rRNA genes from its gff file. We converted the filtered VCF files into consensus sequences using bcftools consensus^221^ with NCTC8163 as a base, masking with N in each different file the positions and regions that do not pass the filtering thresholds.

The genomes in FASTA format were concatenated and an initial ML tree was built using *FastTree*^222^. This tree and alignment was used to detect recombination tracks with ClonalFrameML^143^, using default parameters. This detected 9,369 recombination events across the 170 samples, accounting for a total of 1,241,692 bp in recombination. We then merged the 170 high-quality VCF files with bcftools and used vcftools^216^ to filter out positions missing in more than 25% of the samples, SNPs 5 bp apart, non-variant positions and recombination tracks (*--max-missing 0.75 --mac 1 --thin 5 --exclude-bed recombination.bed).* This resulted in a high quality 6,809 SNPs dataset. We used this dataset to extract positions in the modern *E. tonsillarum* and *Es2* genomes (same GATK and filtering parameters), and the intermediate coverage (between 4× and 1×) *E. rhusiopathiae* genomes (C89, F1364-1436, EDI001,EDI012, ldo50 - minimum depth of 1×).

We used this SNP dataset to build a ML phylogenetic tree using raxml-ng, with the GTR+G model and 1,000 bootstraps. Node support was tested using TBE and FBP. We generated pseudohaploid callings of the SNPs present in this dataset for low coverage samples ((between 1× and 0.05×) using PileupCaller, and projected them in the ML tree using epa-ng. Samples with a first placement LWR over 70% were grafted using gappa^223^. The resultant tree including both projected and non-projected genomes was rooted to the midpoint and visualised using ggtree, collapsing Clades non-relevant for the study. A second tree displaying all clades and all possible projection placements was generated using FigTree^224^.

### Analysis of functional elements and virulence factors

We used a mapping quality filter of 1 to check for the presence of previously identified virulence genes on the *Yersinia pestis* chromosome, pCD1, pMT1 and pPCP1 plasmids, and analysed the coverage of these genes with BEDTools ^225^ v2.29.255. We then visualized the results by making a heatmap using the ggplot2 package in R. As in previous studies (Swaali 2023, Spyrou 2019), we then used IGV to manually inspect the ureD, pde-2 and flhD genes to identify the presence of indels and SNPs specific to these genes, to assess whether they were functional or not.

For *Erysipelothrix rhusiopathiae,* we used the same method to screen a set of known virulence genes ^133,144^. A PCA of the virulence genes was created by building a SNP dataset of those regions in 170 modern and ancient high quality genomes (Clade 1, 2, 3, Intermedia and Ancient). With vcftools we applied a minimum coverage of 5× per position, filtering for and allele depth of 90%, a minimum allele frequency of 0.05, no indels, minimum missingness in the whole dataset of 25% of the samples and minimum genotyping quality of 20 *(--remove-indels --minDP 5 --maf 0.05 --bed virulence.bed --max-missing 0.75 --minGQ 20*). This dataset contained 5,953 SNPs. We created pseudohaploid callings of these positions in lower coverage genomes (5× to 0.05×). We converted the resultant dataset into eig format using convertF, and performed a Principal Component Analysis using smart pca, projection the low coverage data (*lsqproject: YES* and *autoshrink: YES)*^226^. Since most of the ancient samples present genetic affinity against Clade 2, we performed a wide genome Fst analysis (discarding t/rRNA genes) in Ancient (Medieval and Prehistoric) genomes versus Modern Clade 2 genomes. We used the Weir and Cockerham estimator available at vcftools ^227^. We visualised the results using R and selected the top 1% genes with high Fst.

For the discovery of genes with different coverage between modern and ancient strains, we calculated genome wide normalised coverage for every genome. We considered that a gene was present in ancient *E. rhusiopathiae* genomes (>1✕) belonging to human individuals if at least 75% of individuals per group (Early Modern, Medieval, Neolithic-Bronze Age) presented a breadth of coverage of 75% or more per mentioned gene. If more than 75% of individuals presented less than 75% of breadth of coverage, we considered the gene as absent.

### Radiocarbon dating

All dates from human bone previously published in Iaia and Dolfini^53^ were calibrated with the IntCal20 atmospheric curve^54^ in OxCal v4.4.4^55^. The dates were modelled as a single phase since they were derived from bones within the same stratigraphic layer (US 8), and dates were rounded out to 10 years^228^ [Supplementary Figure S10].

### Isotopic analysis

A sample of the mandibular cortical bone from individual GSP005 was measured for bulk δ^13^C and δ^15^N isotopic analysis, within a dataset of 30 samples with acceptable collagen yields from the *Grotta della Spinosa*^229^. The cortical bone may represent an average intake of protein sources within a range of approximately the last decade or so of an individual’s life^230^.

## Availability of data and material

The datasets generated and analyzed during the current study are available in the ENA repository under the accession ID: PRJEB108536 (Individuals from Grotta della Spinosa - GSP) and PRJEB108537 (Cova Bonica - CB13).

## Acknowledgements

The authors are grateful to Dr. Enrico Maria Giuffrè (SABAP Siena, Grosseto and Arezzo) for permission to analyse the *Grotta della Spinosa* human remains, and to Dr. Antonietta del Bove for providing information about the assemblage. This work was supported by the European Union through the European Research Council Advanced Grant ‘Making Ancestors: The Politics of Death in European Prehistory’ (No. 885137) (TdD, CLS, BB, JER, MAT, JT, SP, HK, TS), UKRI Engineering and Physical Sciences Research Council Horizon Guarantee (EP/Y009878/1) (CLS, BB), Estonian Research Council grant PUT (PRG1899) (TdD, MT, HK, MM), Estonian Research Council grant PUT (PRG1027) (BB, HK, TK), European Union and Estonian Research Council through the Mobilitas 3.0 (MOB3JD1225) (RB), Ministry of Education and Research Centres of Excellence grant TK215 name of CoE (SS).

## Author Information

### Authors’ contributions

Conceptualisation – TdD

Formal analysis – TdD, BB, JET, MAT, TS

Funding acquisition – CLS, JER, MAT

Investigation – JET, TS, SP, HK, BB, SS, SB, AS, MT

Methodology – TdD, BB, JET, SP, TS

Resources – MG, MT, BA,TK, CLF, JD, MS, RB, CLS, MM

Supervision – CLS, MAT, JER

Visualisation – TdD, BB, JET, SP, BA, TS

Writing – original draft – TdD, BB

Writing – review & editing – All authors

## Ethics declarations

### Ethics statement

All necessary permits were obtained for this study, which complied with all relevant regulations. The Soprintendenza Archeologia, Belle Arti e Paesaggio per Siena, Grosseto e Arezzo authorised this research to be carried out on the human remains from *Grotta della Spinosa* (stored at the Museo di Antropologia Giuseppe Sergi, University of Rome Sapienza). This research received approval from the Ethics Committee of the Department of Archaeology, University of Cambridge and followed internal project guidelines, including the full recording of remains prior to removing samples. Data analyses were carried out with the facilities of the High-Performance Computing Center of the University of Tartu. In an effort to obtain multiple biomolecular results on the same elements, where possible, teeth for aDNA analysis were selected from elements previously subjected to bulk δ^13^C and δ^15^N isotope analysis (Bernardini 2022).

## Additional information

Supplementary Information is available for this paper.

