## Supplementary Information for "Co-occurrence of Yersinia pestis and other zoonoses during European prehistory"

### SI1 - Selection of archaeological samples

The human remains from Grotta della Spinosa were transferred to the Museo Giuseppe Sergi at the University of Rome Sapienza with the permission of the Superintendency. They were previously studied to estimate the minimum number of individuals[1], and included in a research project examining Copper Age dietary and mobility practices through stable isotopic analyses[2]. The full osteological assemblage (26,800 fragments) was re-analysed by two of the authors (JET and SP) as part of the ERC-funded 'Ancestors' Project. All human bone fragments were inventoried, identified to element, side, and part of element when possible, and their taphonomic condition and bioarchaeological profile (age, sex, health and pathology) were recorded.

Given the commingled depositional context of the human remains and the very large number of fragments in the assemblage, elements for aDNA sequencing were carefully selected according to several criteria. The petrous portion of the temporal bone (*pars petrosa*) was sampled (n=10) according to the side which was most numerous in the assemblage, in an effort to prevent double sampling. Dentition (n=13) was mostly retrieved from mandibular alveoli, to allow for observations regarding dental health, pathology and osteological sex estimation. The majority of samples was selected according to the most numerous side and tooth (lower left first molar), although some samples were from alternate teeth where we could be reasonably certain these pertained to different individuals. Owing to the extent of fragmentation, most teeth did not have antimeres present in the same mandibular fragments. Each tooth was fully recorded and photographed and efforts were made to minimise the sample size. Additionally, teeth were preferentially selected from mandibles which had previously been sampled for bulk isotopic analysis[2] and which had calculus deposits for proteomic and microbial aDNA analyses, enabling multiple biomolecular and geochemical observations to be retrieved from the same element.

The sample reported here (GSP013) was obtained from a lower left first molar extracted from a fragmentary mandible of an adolescent. The mandible was less than 50% complete and broken in two fragments which were missing the central symphyseal portion of the element. The quality of the cortical bone is poor and mid-brown soil was adhered to much of the bone surfaces. The element had been broken post-depositionally, with breakage to mineralised bone observed, and it had been sawn on the internal aspect of the ramus on the right side to remove a bone sample for bulk isotope analyses (see below)[2]. The absent portion of the element pertained to tooth positions for the lower incisors and canines; as a result, alveoli were present and visible bilaterally for positions from the first premolar to the third molar. However, only eight teeth were present, as the lower left second and third molars were missing post-mortem. The right third molar was observed to be erupting and root development was approximately half-complete. The roots of both first molars presented half closed apices, while root development for the second molars was not observable. There was no evidence of periodontal disease or periapical lesions and only pinprick-sized areas of dentine were exposed on some molar cusps. Dental development for this individual estimates their age at 17±1 years[3].

**SI2 - Paleopathological analysis of GSP013**

Consistent with their young age, their dental and alveolar health appeared to be good at the time of death and the extant mandibular dentition displayed minimal attritional wear. The first molars and right second molar presented linear enamel hypoplastic (LEH) lesions, which indicate growth disruptions during the formation of the enamel and may be attributed to general physiological, pathological, environmental or nutritional stressors during relatively short periods of time[4,5]. Two lesions on the lower left first molar were estimated to have occurred between the ages of 2.09–2.17 years, and one lesion on the lower right first molar shortly after, around 2.62 years[6]; a further lesion on the lower right second molar indicated a later disruption, around the age of 5.91 years. Some dentitions are missing, notably the canines and the lower left second molar, which represent long durations of development in crown formation (from circa 0.5–5.5 years and 2–8.5 years, respectively, see[7]). With only one LEH observed on the extant right second molar, the apparently minimal evidence growth disruptions in their later childhood may suggest that this individual was infected closer to their time of death, or at least after approximately 7 years of age, which is the latest age at which LEH lesions are usually estimated to form[4].

Isotopic measurements from the adolescent considered here are within the range of values observed from the site overall (Table S4). This indicates that their diet was similar to that of the wider burial group, which was predominantly based on terrestrial animals and dairy products, with a lesser contribution of $C_3$ plants, and likely included very occasional freshwater resources. Bulk isotopic data are coarse-grained, however, and we cannot exclude the possibility that this individual's diet may have changed toward the end of their life.

**SI3 - *Yersinia pestis* gene analysis**

We manually explored *ureD*, *PDE-2* and *flhD*. The *ureD* gene (urease D), encodes an essential component for urease activity, which catalyzes the conversion of urea to ammonia and carbon dioxide. In *Y. pseudotuberculosis*, this activity supports survival under acidic conditions. In *Y. pestis*, however, the production of ammonia is toxic to the flea midgut, and thus, *ureD* has been inactivated during evolution. A six-guanine (poly-G) insertion at position 2,997,296 in strain CO92 disrupts the gene's function, eliminating urease activity and aiding flea adaptation[8]. *flhD*, part of the *flhDC* operon, is a regulator of flagellar biosynthesis. In *Y. pseudotuberculosis*, *flhD* expressed in a temperature dependent manner. However, all analysed *Y. pestis* strains harbor a frameshift mutation that disrupts *flhD*, resulting in loss of motility[9]. *PDE-2* regulates cyclic-di-GMP turnover, impacting biofilm degradation. In *Y. pestis*, biofilm formation is essential for blocking the flea's proventriculus, which enhances transmission efficiency. A T insertion at position 1,434,044 in the *PDE-2* coding sequence causes a frameshift that renders the gene inactive[10].

### SI4 - Ancient *Erysipelothrix* screening and validation

We used the AncientMetagenomeDir[11] to select historical and prehistoric raw data coming from individuals infected with *Yersinia pestis.* We additionally included individuals infected with *Salmonella enterica,* and those of domestic species suspected of carrying the bacteria. We also included 2,572 samples from other 79 published studies.

After preprocessing the samples using AdapterRemoval2[12] with the same parameters used for ancient *Y. pestis,* we proceeded to screen against those selected individuals by mapping against a set of *Eryspelothrix spp.* assemblies. We used *bwa* backtrack, identifying as positive for a species sample when a sample reached a depth of coverage above 0.01×, breadth of 1%, average edit distance under 1.5, and decaying edit distance values to the mapped species. We also analysed terminal deamination patterns[13], PMD score [14]distribution and read length distribution. Final species assignment was validated using phylogenetic analysis.

Due to its positioning, C10091 and ADN004 were realigned using a competitive mapping against the different *Eryspelothrix* reference genomes, and non *E. tonsillarum* sequences were reclassified using blast[15]. Mapped sequences were screened using blastn against the core_nt database. Results were classified to the LCA using MEGAN6 blast2lca[16]. Sequences were considered to be classified in a taxonomic level if 99% of the sequences belong to a taxonomic level under it. The resultant screens reveal the presence of *Eryspelothrix-like* sequences in both samples that cannot be classified. Those 2 samples were subsequently discarded from downstream analysis.

We used PROKKA to annotate the assemblies of 18 genomes of *E. rhusiopathiae* and 2 of *E. tonsillarum*[17]. Roary was used to account for the species divergence using a *minimum* percentage identity for blastp of 90[18]. A total of 636 core genes were identified, which were used to map 196 modern genomes of *E. rhusiopathiae* and 12 modern genomes of *E. tonsillarum* using *bwa mem*[19]. We used the same set of genes to map the retrieved ancient *E. rhusipatiae* and *E. tonsillarum* genomes, using bwa backtrack with parameters adjusted for aDNA data (-l 10,000 -n 0.01 -o 2)[20,21] against the *E. rhusiopathiae* reference genome NCTC8163. Following this, we used GATK[22] Unified genotyper to create variant callings of the mapped genes. We filtered variants using vcftools[23]. For modern samples we considered a minimum depth of 5, allele depth ratio of 0.9, minGQ of 20, and no 'heterozygous' positions. For ancient samples, genomes with a depth ≥4× (Galicia3, 6Post, KIL034, LSC005A, EDI012, HTC009, FRA012, and CB13) were filtered using the same parameters. This initial dataset was then merged using bcftools[24], and filtered again using vcftools, this time keeping variant positions covered in at least 99% of the samples, and keeping only biallelic positions. This high quality dataset contained 40,629 biallelic positions. For ancient samples with a depth ranging from 5× to 1× (ldo50, ED124, C89, F1364-1436, EDI001), positions in the high quality dataset were called using *GATK Unified Genotyper*[22] , and with same filtering parameters as before, with a minimum depth of coverage of 1×. Finally, low quality (<1×) and *E. tonsillarum* sample variants were called by selecting a random allele present in the data using PileupCaller[25].

An exploratory phylogeny to confirm the detected species was built using modern, high quality and medium quality genomes. We used raxml-ng[26] with a GTR-G model and 100 bootstraps. Low quality genomes were projected using epa-ng, and grafted using gappa. The resultant tree was rooted to the midpoint and displayed using *ggtree*[27].

In order to avoid erroneous mapping of sequences belonging to *Yersinia pestis* and *Salmonella enterica* in coinfected individuals. We have followed two approaches. The first one was to assess the affinity of the *Y. pestis* and *S. enterica* via mapping. We started by converting the *Y. pestis CO92* and the *S. entrica Paratyphi C RKS4594* reference genome into 100bp fastq reads. We then used *bwa backtrack* with the same parameters used in ancient samples (-n 0.01, -l 10,000 -o 2)[20,21] to map those sequences against the *E. rhusiopathiae* reference genome. After keeping mapped sequences with a mapping quality equal or above 0, we have calculated basic statistics with Qualimap2[28], and read distribution across the reference genome (10,000bp windows) using bedtools.

### SI5 - Differences on phylogenetic placement of *Ersypelothrix rhusiopathiae* retrieved from shotgun data versus sequences retrieved from capture

For EDI001, we have retrieved a total of 467 *Ersypelothrix rhusiopathiae* sequences from *Y. pestis* capture data, and 47,575 from a tooth-root shotgun library (both mapping quality 30 sequences without duplication). We have independently projected both groups of sequences into a *E.rhusiopathiae* ML phylogeny using epa-ng [Figure S14][29] The tree was created using 6,908 high quality recombination free SNPs found across 196 modern *E. rhusiopathiae* genomes. Clades 1 and 3 are collapsed, displaying only Clade 2 (average TBE value of 99). Shotgun derived sequences are positioned near other Early Medieval genomes (ldo050) with a high first positioning likelihood weight ratio (fpLWR) of 99% among other Clade 2 genomes[30]. Sequences derived from *Y. pestis* capture library are projected in several positions with low fpLWR (~9%) spread across Clade 2. Both libraries do not cluster together. We conclude that although not highly accurate, *E. rhusiopathiae* sequences recovered from *Y. pestis* capture libraries are valuable to have a tentative positioning of the genome and to which potential Clade does the genome belongs to. We could not compare other individuals since the amount of sequences in *Y.pestis* capture versus shotgun is otherwise negligible in FRA005 (16 versus 7,850), GSP013A (241 versus 17,663), and C10091 (0 versus 7,711).

### SI6 - Comparison of *Ersypelothrix rhusiopathiae* sequences obtained from multiple tissues and libraries

We have compared *E rhusiopathiae* presence in different libraries from the same individuals' different tissues. For GSP013 - 005, we have data from 2 libraries obtained from the left petrous bone (GSP005A and B), and a library from the first left lower molar (GSP013A). For the petrous bone libraries we found a total of 64 mapping against *E. rhusiopathiae* (54 and 10 reads respectively), while the tooth root presents 17,663 sequences. For the other individual that we have libraries from different tissues available, EDI001 (maxilla, dentin powder and tooth root) we find that again the tooth root is concentrating the bulk of *Erysipelothrix* with a total of  48,689 of sequences, against 66 in the dentin powder, and 43 in the maxilla. These results are to be expected from a pathogen with septic potential, where bacteria would be found in the highly vascularised pulp chamber, in comparison to less vascularised skeletal elements such as petrous bone or  postcranial elements[31]. This behaviour has been observed in *Yersinia pestis* bubonic plague[32,33], and *Salmonella enterica* paratyphoid serovars in which enteric disease can derive into blood infection[34,35], or *Plasmodium* parasites, which have life stages invading erythrocytes [36,37]. Other pathogens with non-strict infectious stages present in blood can be found in teeth, including *M. leptrae,*[38] *T. pallidum* [39] can be retrieved.

Those preliminary results here suggest that the presence of the bacteria only in tooth roots, but not other skeletal elements of EDI001 and GSP013, could be attributed to a septicemic form of  *E. rhusiopathiae* infection. This bacteria presents 3 main infectious manifestations. A contained skin infection, usually in the form of a rash[40], a more severe diffuse skin infection[40], and a third form (more rare) involving septicemia (specially in immunocompromised individuals) [41–45] and fatal endocarditis[40]. The presence of bacteria in blood during this last manifestation could be responsible for the presence of bacterial sequences in tooth roots. Another scenario that we can not rule out is the stochasticity associated with pathogen aDNA retrieval, in which different elements or even extractions do not present the same levels of pathogen DNA< making identification difficult[32,46].

### Supplementary Figures

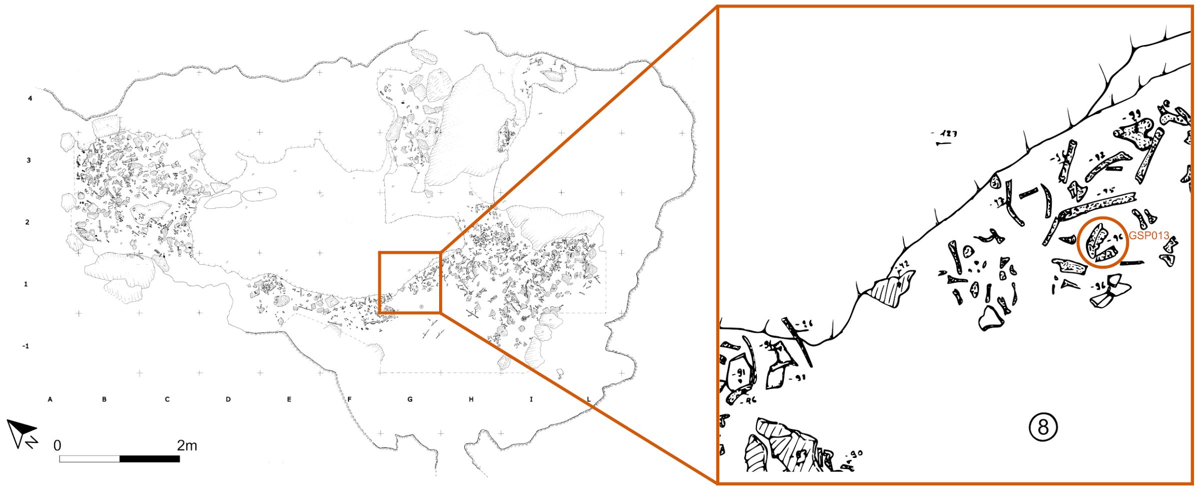


**Supplementary Figure S1. Diagram of Grotta della Spinosa.** Plan of human bone deposits in layer 8 level 3 of Grotta della Spinosa (scale: 1:10), showing the position of the mandible belonging to sample GSP013. Redrawn after R. Guidi (made with Inkscape v. 1.3.2). Permission to reuse the image granted by Dr. Biancamaria Aranguren.

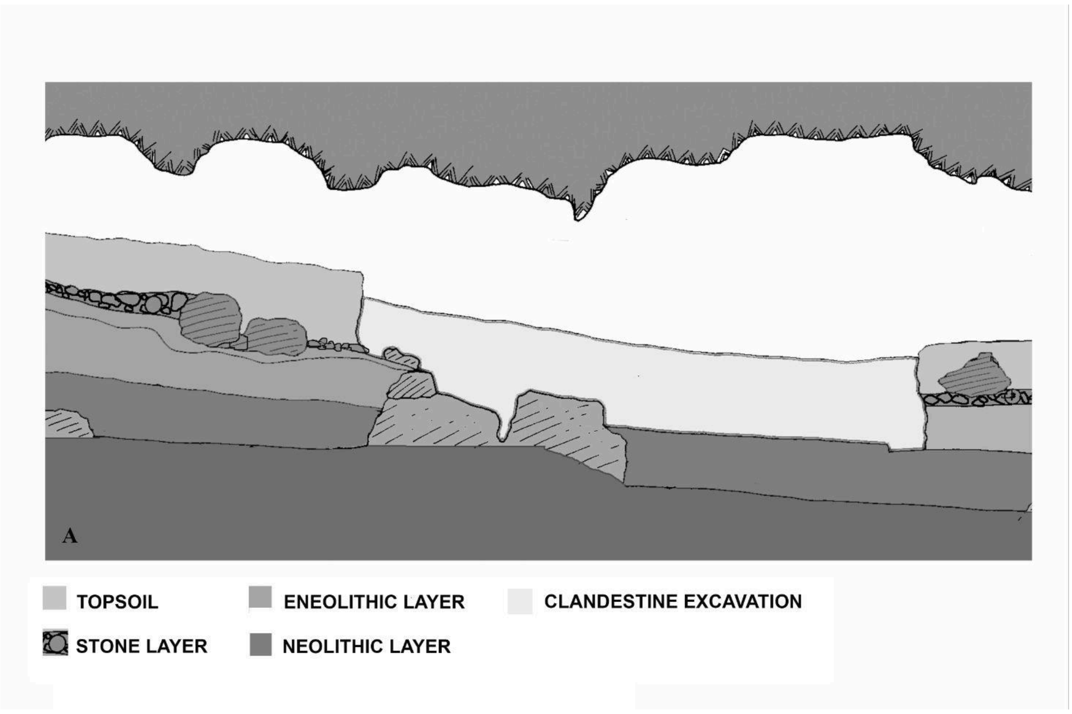


**Supplementary Figure S2. Archaeological context of *Grotta della Spinosa*. A** Sagittal section of the site.

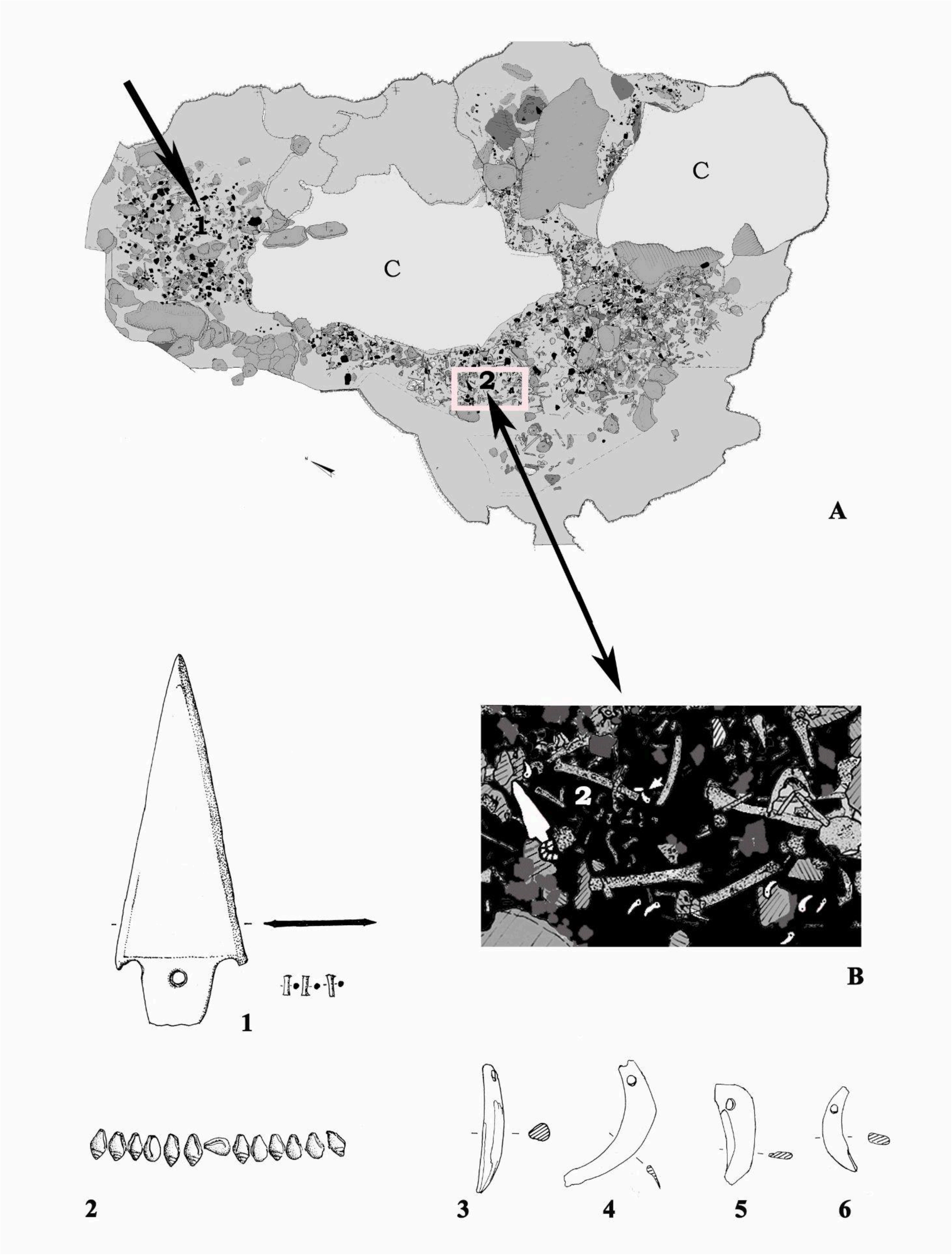


**Supplementary Figure S3. Location of the human remains and artifacts within the cave.** Permission to reuse the image granted by Dr. Biancamaria Aranguren.

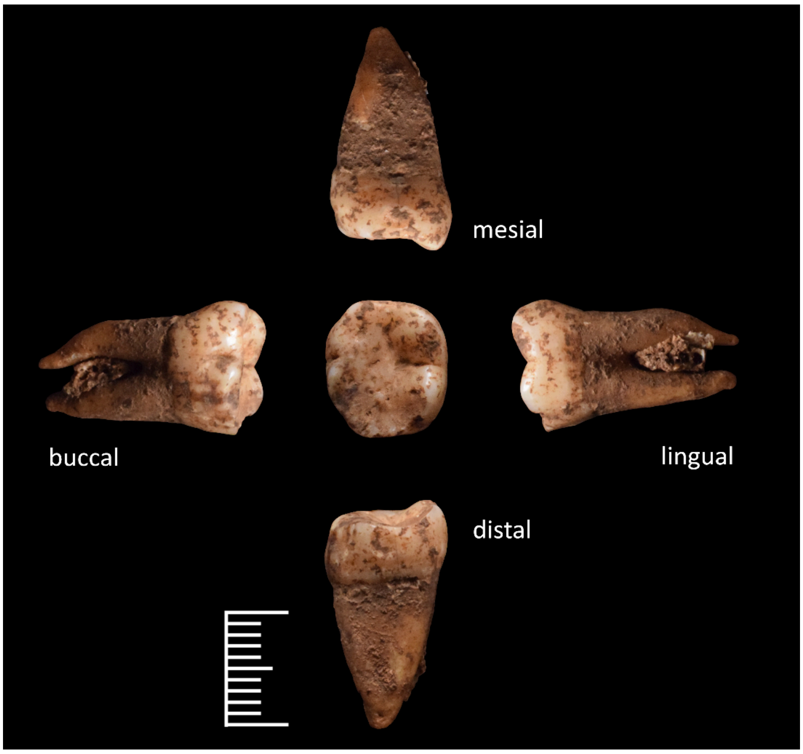


**Supplementary Figure S4. GSP013 tooth sample.** All aspects of the lower left first molar (GSP013) from mandible in layer 8, level 3, quadrant G1, prior to sampling; scale bar: 1 cm.

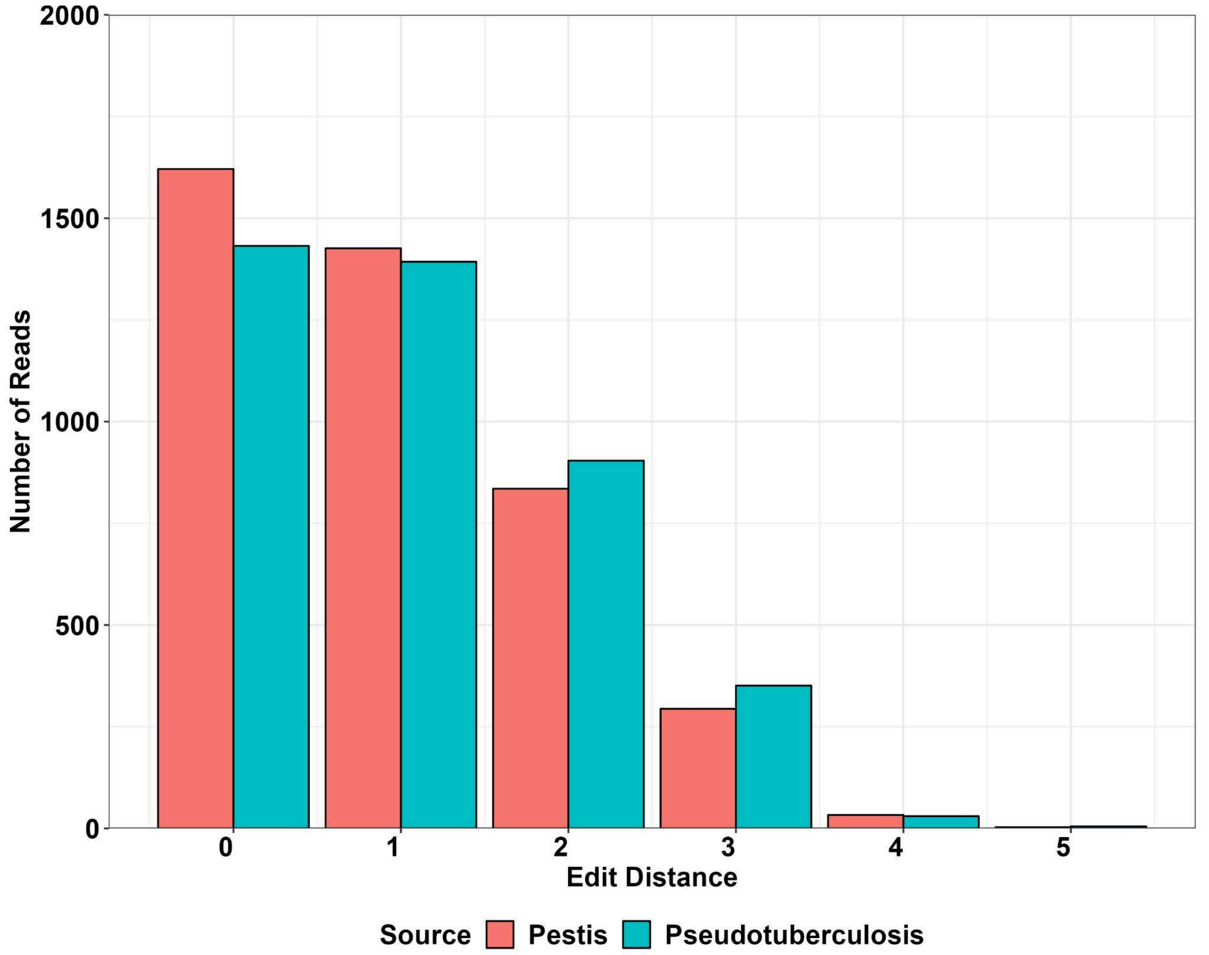


**Supplementary Figure S5. Reference affinity of GSP013.** Edit distance of GSP013 sequences mapped against *Y. pestis* and *Y. pseudotuberculosis* reference genomes. GSP013 displays more mapped reads and lower edit distances to *Y. pestis*, demonstrating the presence of the bacteria in the screening data.

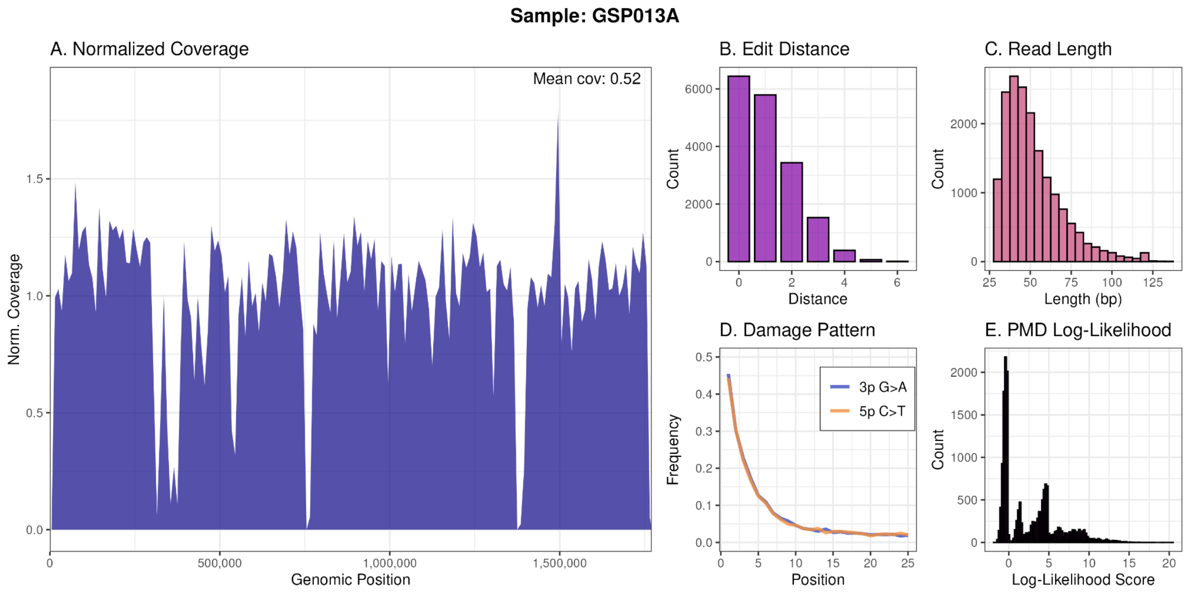


**Supplementary Figure S6. Mapping statistics and damage of *Erysipelothrix rhusiopathiae* retrieved from GSP013.** A) Coverage depth across the genome. B) Edit distance distribution. C) Read length distribution. D) Frequency of 3’ G to A and 5’ C to T substitutions along the reads’ last 25 basepairs. E) Mapped reads’ PMDscore distribution.

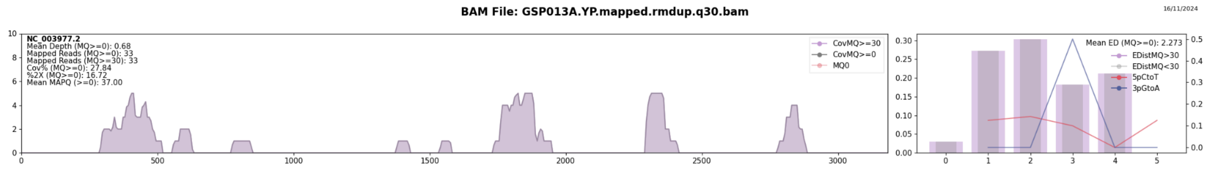


**Supplementary Figure S7. Mapping statistics and damage of Hepatitis B virus retrieved from GSP013.**

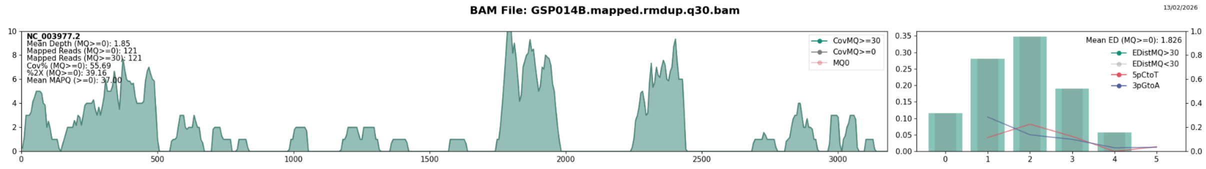


**Supplementary Figure S8. Mapping statistics and damage of *Hepatitis B virus* retrieved in GSP014.**

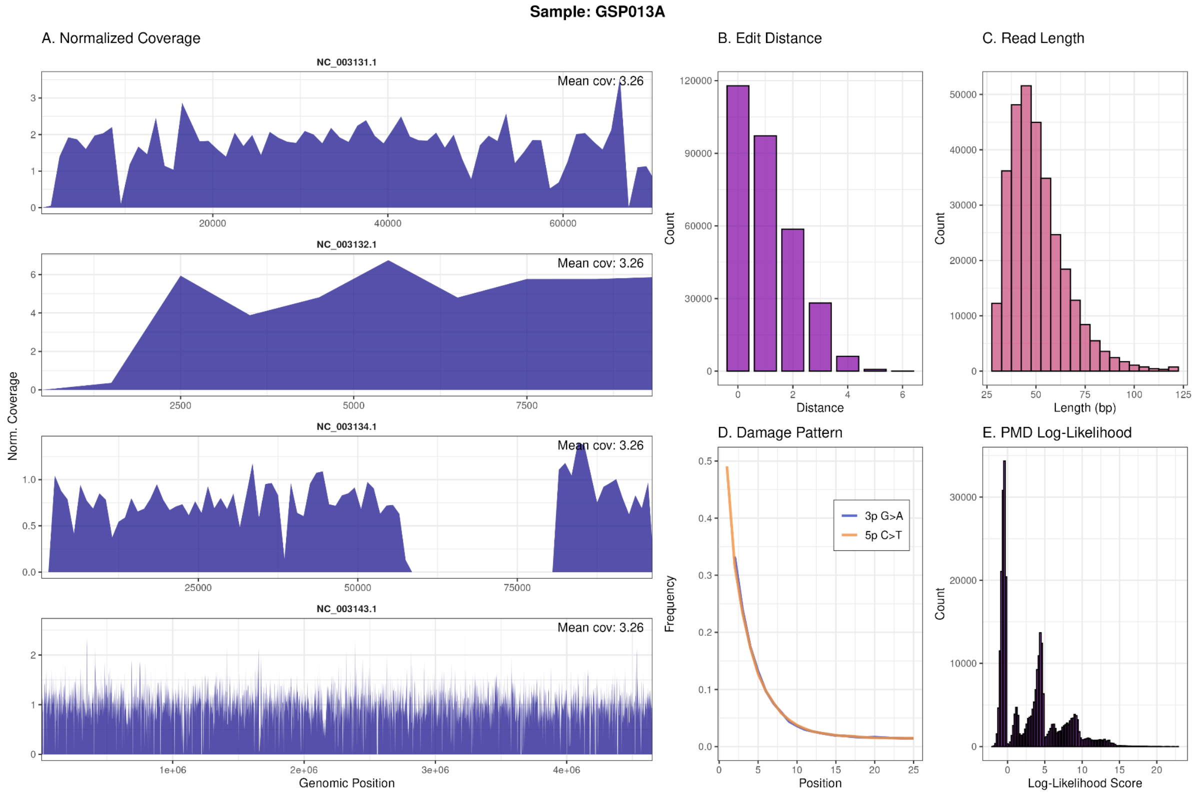


**Supplementary Figure S9. Mapping statistics and damage of *Yersinia pestis* retrieved in GSP013 (Post-capture).** A) Coverage depth across the genome. B) Edit distance distribution. C) Read length distribution. D) Frequency of 3' G to A and 5' C to T substitutions along the reads' last 25 basepairs. E) Mapped reads' PMDscore distribution.

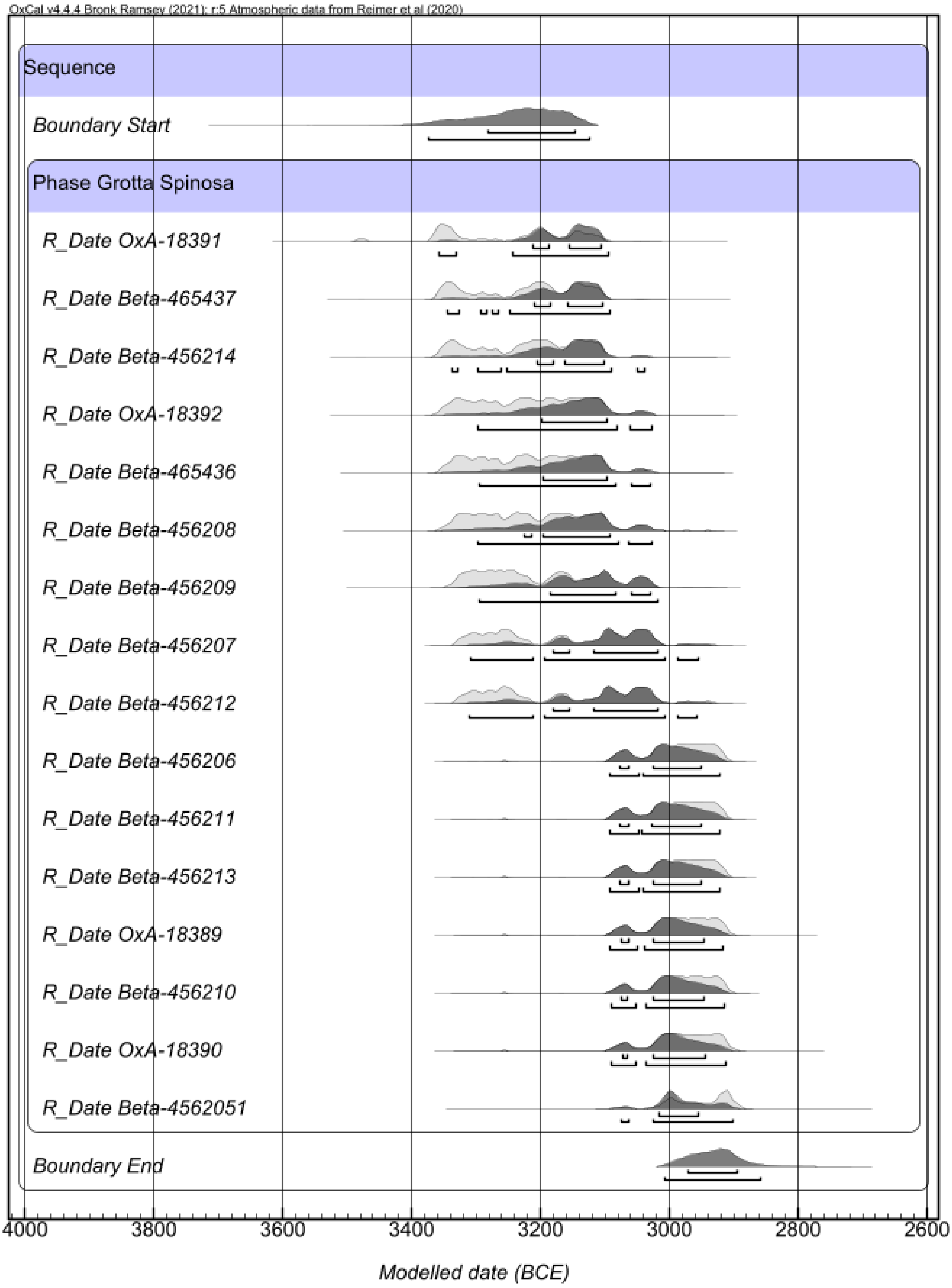


**Supplementary Figure S10. OxCal plot of the Bayesian model of all radiocarbon dates from human bone at Grotta della Spinosa.** The calibrated range of each date is shown in light grey, with the range modelled on the posterior density estimate in dark grey.

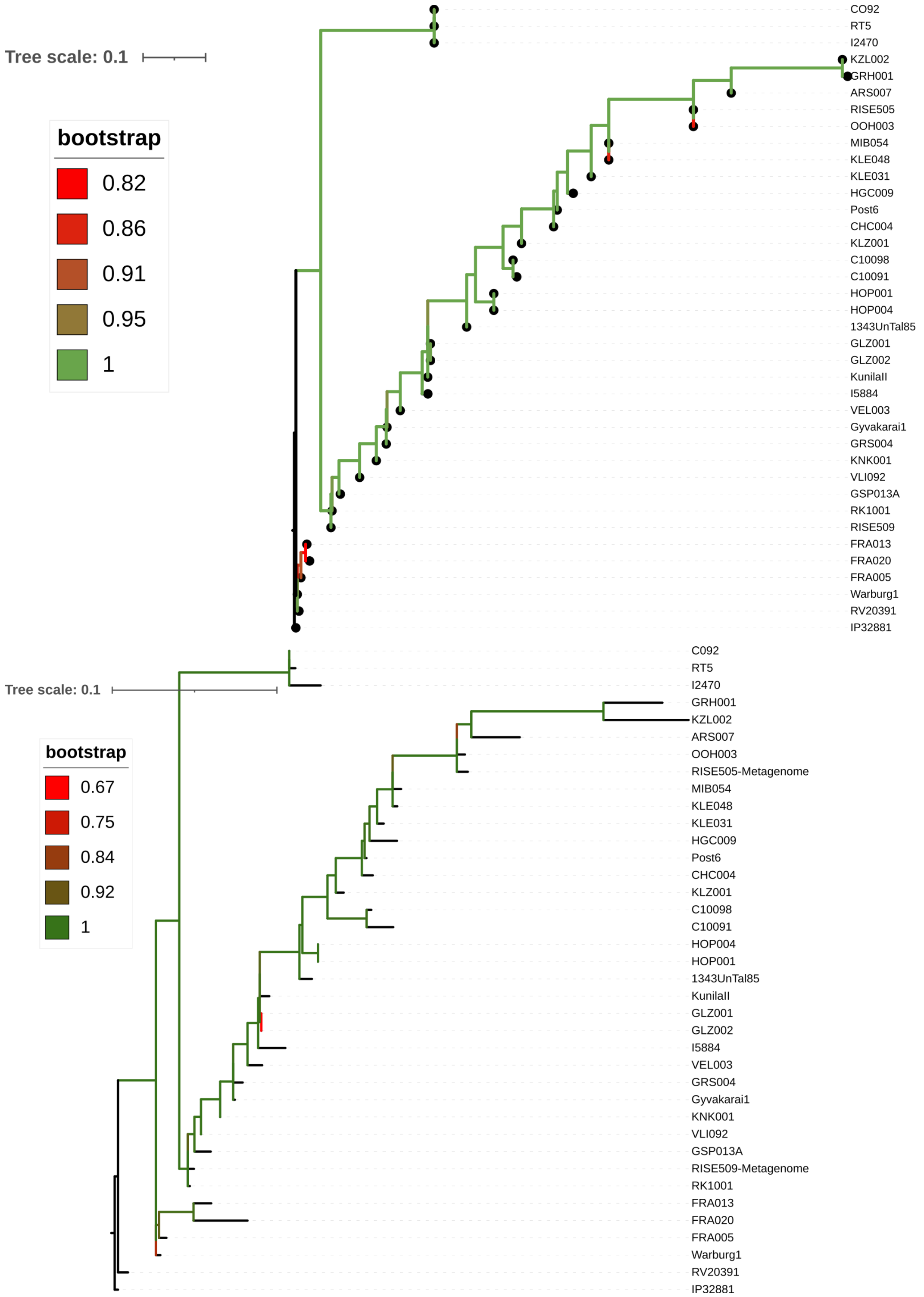


**Supplementary Figure S11. *Y. pestis* ML trees of GSP013A with other modern and prehistoric genomes.** Tree without singletons (top) and without C ↔ T and G ↔ A transitions (bottom) used to assess effect of errors due to low coverage and aDNA damage. Bootstrap values (TBE) are displayed using a colour gradient.

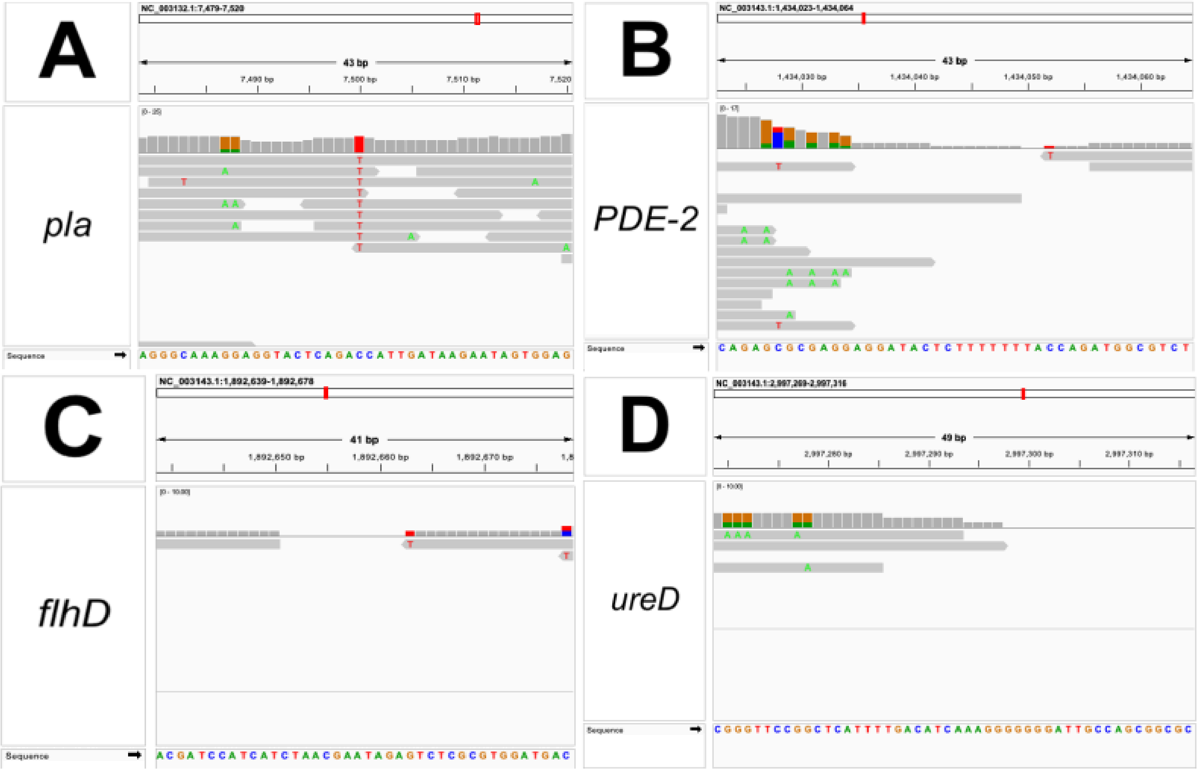


**Supplementary Figure S12. Manual evaluation of genes involved in flea transmission and virulence, using IGV [47,48].** A) We evaluated the pPCP1 plasmid and we observed the ‘T’ variant at position 7,500 of the plasmid. This corresponds to the ancestral variant and results in an isoleucine at amino acid position 259 of the pla gene [49]. B) PDE-2 C) flhD D) ureD.

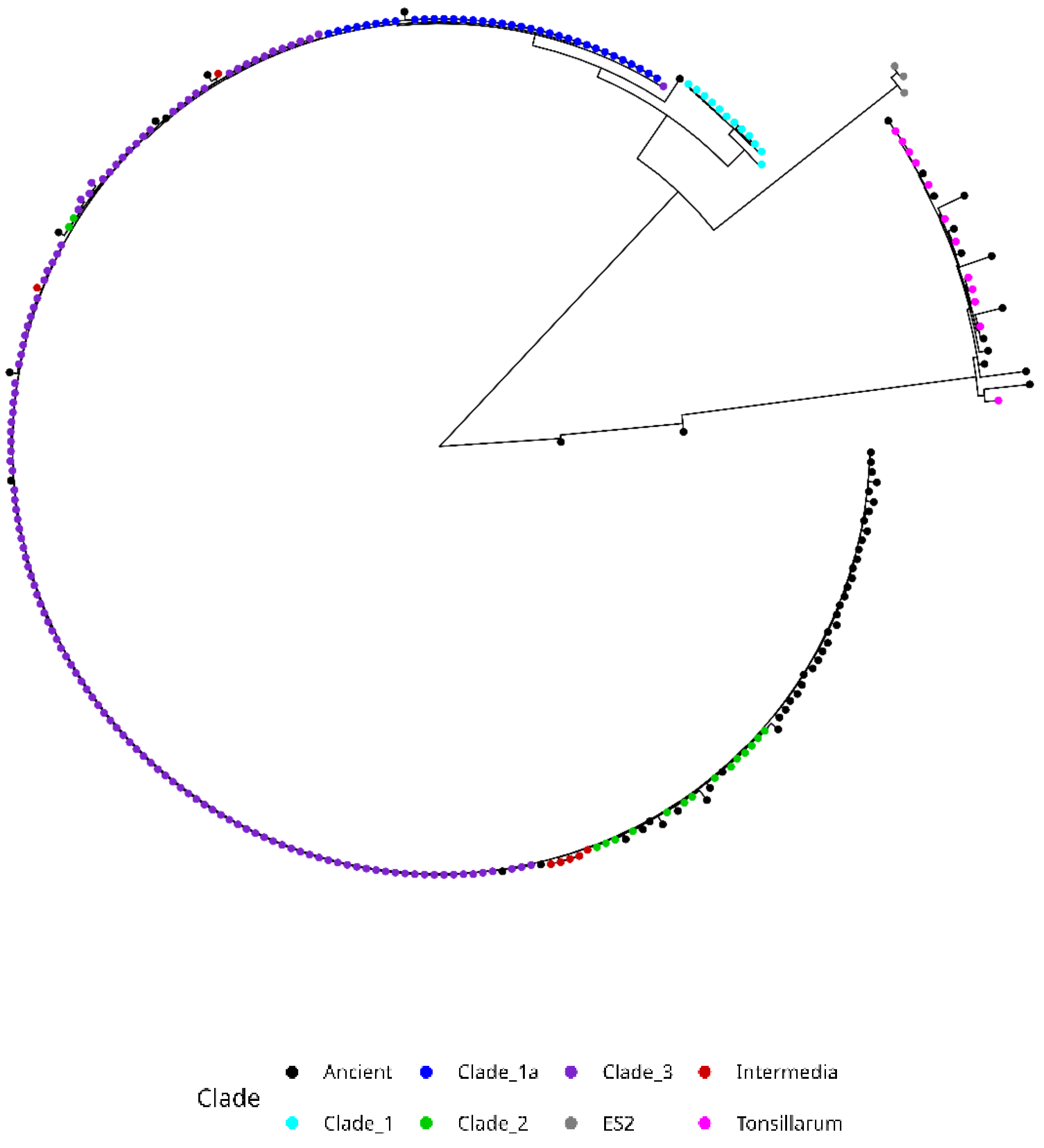


**Supplementary Figure S13.** Exploratory ML (raxml-ng 100 bootstrpas - GTR+G) phylogeny of 196 modern *E.rhusiopathiae* genomes, 3 *E. species S2, 12 E. tonsillarum* and 64 Ancient *Erysipelothrix.* Clades and Ancient status is represented per sample using the tip point colour.

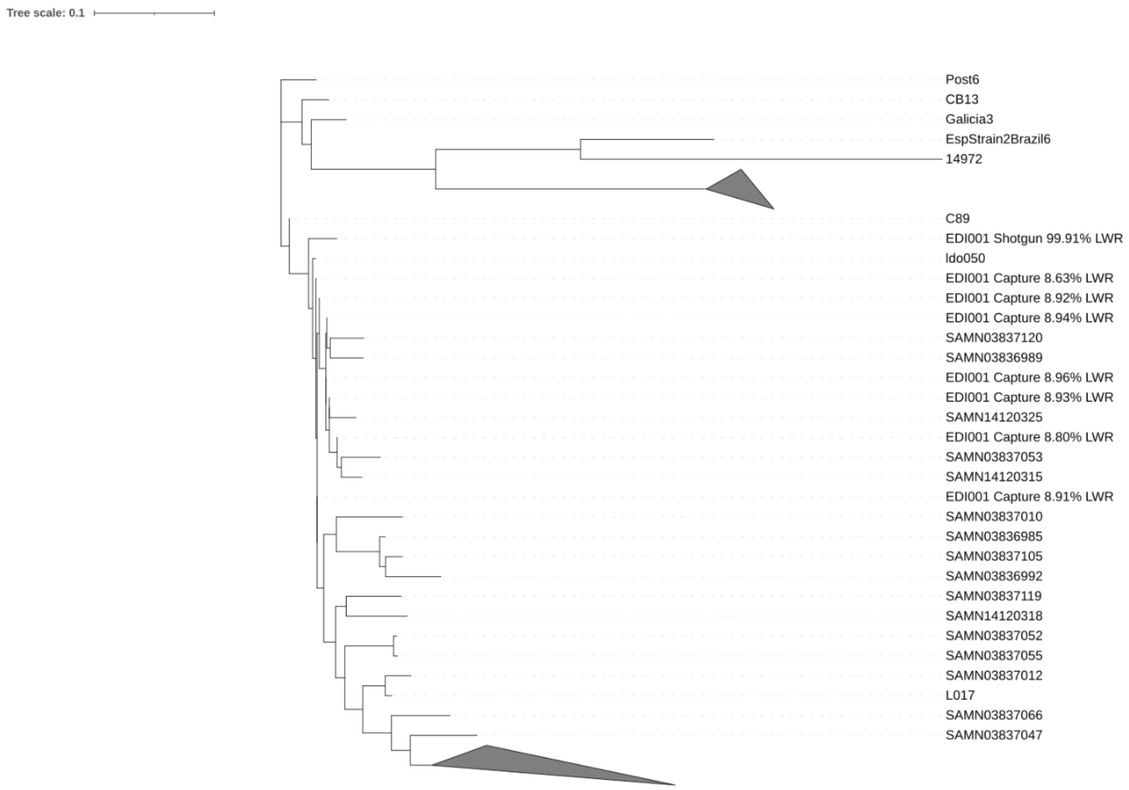


**Supplementary Figure S14.** Comparison of phylogenetic positioning after epa-ng projection of *E. rhusiopathiae* sequences recovered from 2 different libraries (shotgun versus *Y. pestis* capture) of individual EDI001 in the context of modern *E. rhusiopathiae*.

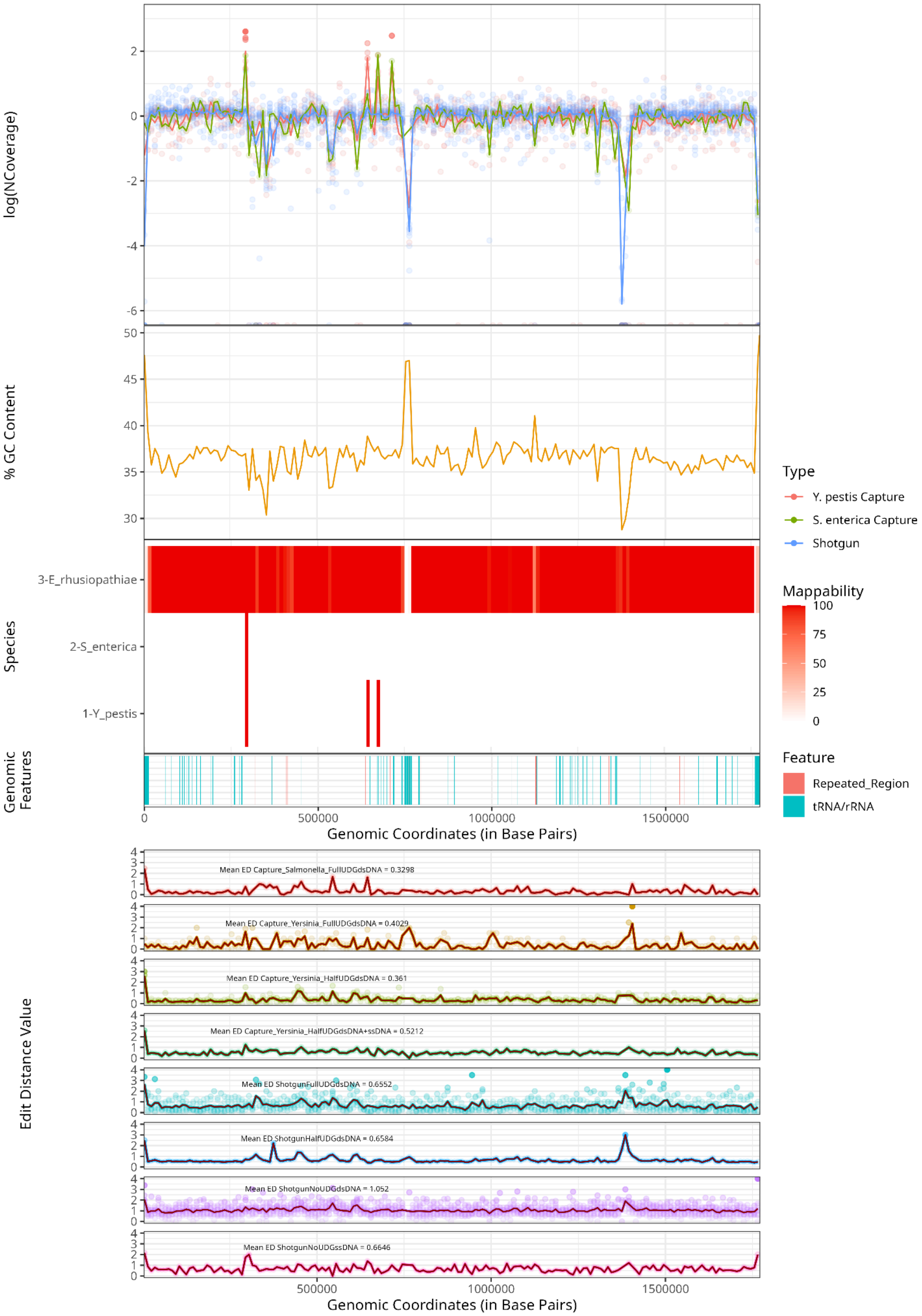

log(NCoverage)
% GC Content
Species
3-E_rhusiopathiae
2-S_enterica
1-Y_pestis
Genomic Features
Genomic Coordinates (in Base Pairs)
Type
Y. pestis Capture
S. enterica Capture
Shotgun
Mappability
Feature
Repeated_Region
tRNA/rRNA
Edit Distance Value
Mean ED Capture_Salmonella_FullUDGdsDNA = 0.3298
Mean ED Capture_Yersinia_FullUDGdsDNA = 0.4029
Mean ED Capture_Yersinia_HalfUDGdsDNA = 0.361
Mean ED Capture_Yersinia_HalfUDGdsDNA+ssDNA = 0.5212
Mean ED ShotgunFullUDGdsDNA = 0.6552
Mean ED ShotgunHalfUDGdsDNA = 0.6584
Mean ED ShotgunNoUDGdsDNA = 1.052
Mean ED ShotgunNoUDGssDNA = 0.6646
Genomic Coordinates (in Base Pairs)

**Supplementary Figure S15.** (1st panel) Log normalised coverage distribution across *E. rhusiopathiae* reference genome. (2nd panel) GC content percentage across *E. rhusiopathiae* reference genome. (3rd panel) *E. rhusiopathiae* mappability and regions with *Y. pestis* and *S. enterica* potentially map. (4th panel) Presence of repetitive regions (>30bp) and t/rRNA across *E. rhusiopathiae* genome.(5th panel). Different in library type edit distance value across the genome.

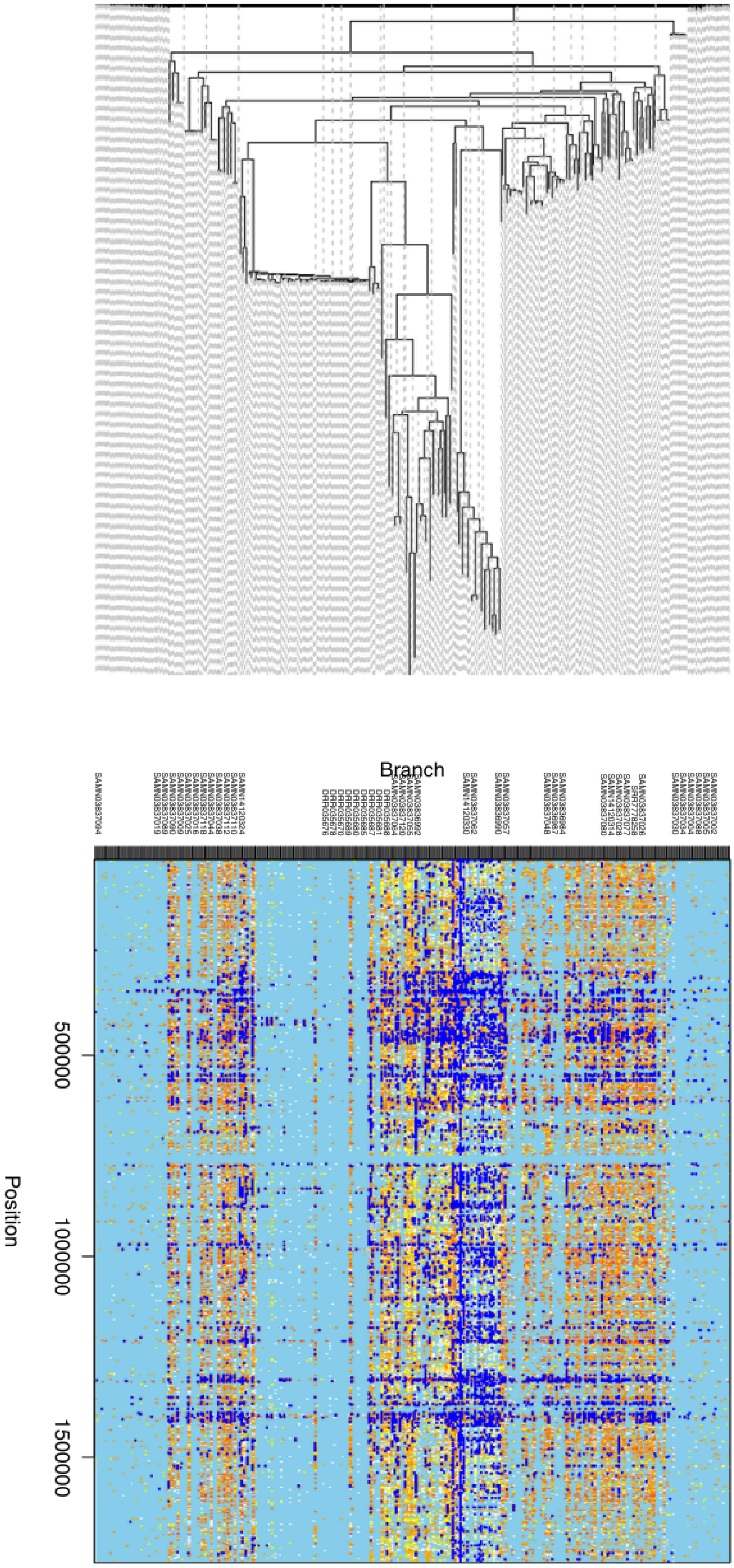
Branch
Position
50000
100000
150000

**Supplementary Figure S16.** Location across *E. rhusiopathiae* reference genome of recombination events detected by ClonalFrameML in modern *E. rhusiopathiae* diversity and 8 high Coverage ancient genomes (Galicia3, CB13, LSC005A, 6Post, FRA012, EDI012, KIL034, and HTC009).

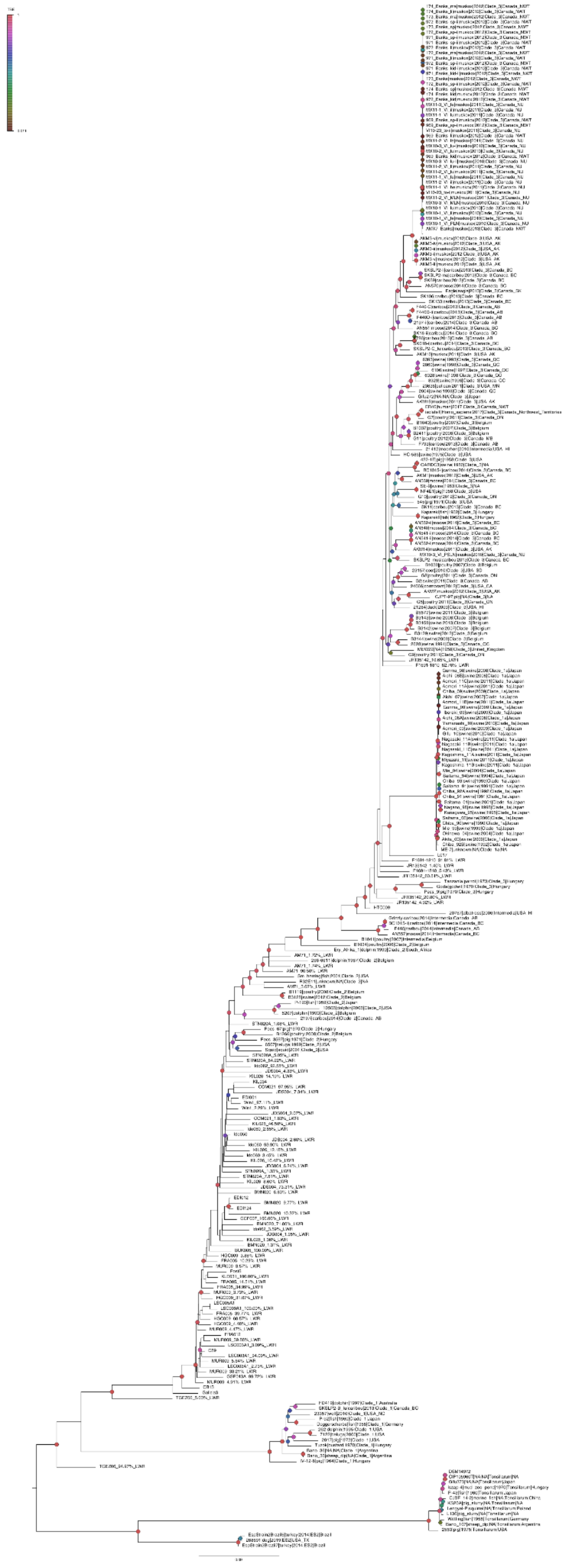

**Supplementary Figure S17.** E. rhusiopathiae ML tree (1,000 bootstraps, GTR+G, 6,809 High-quality recombination-free SNPs, TBE Bootstrap values are displayed using a colour gradient) displaying all epa-ng low-quality genomes projection possibilities with their respective LWR.

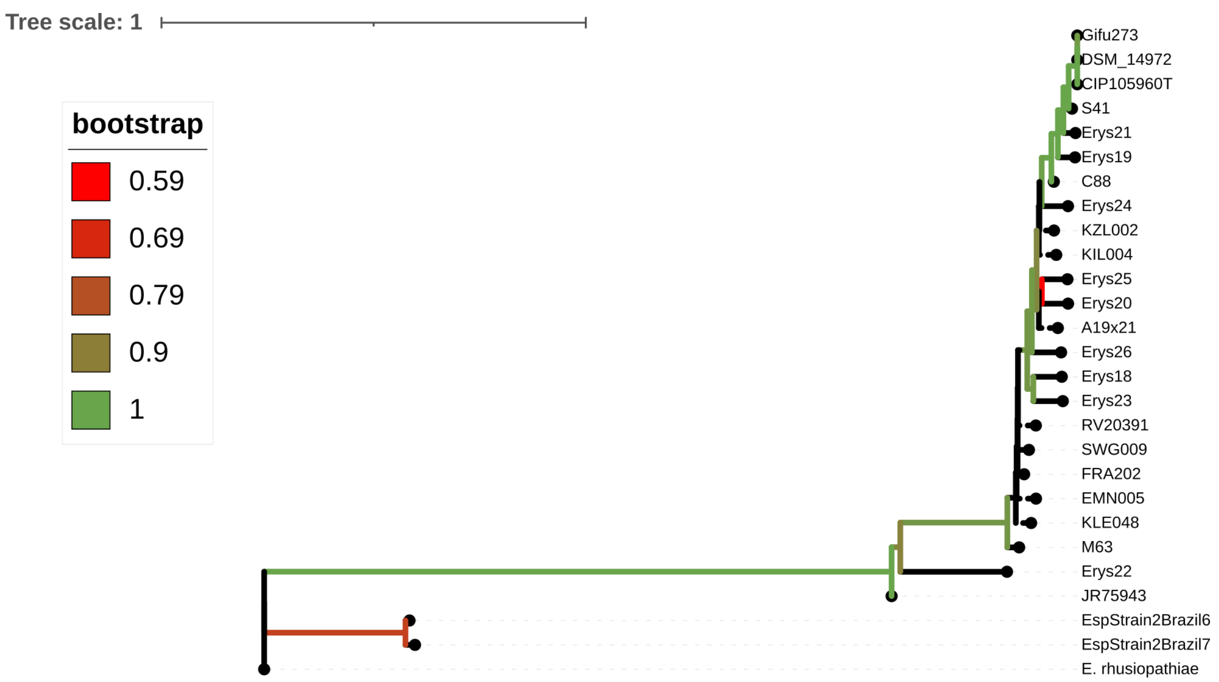


**Supplementary Figure S18.** *E. tonsillarum* and *Es. species* ML phylogeny including 60,509 High-quality recombination-free found across 14 modern strains of the bacteria. Positions in this dataset were called in medium quality genomes (between 4× and 1×; C88, JR75943, M63 and S41). Low quality samples (from 1× to 0.05×) were called with pseudohaploid callings using Pileup caller. The tree was built using the GTR+G model and 1,000. TBE bootstrap value is represented using a colour gradient.

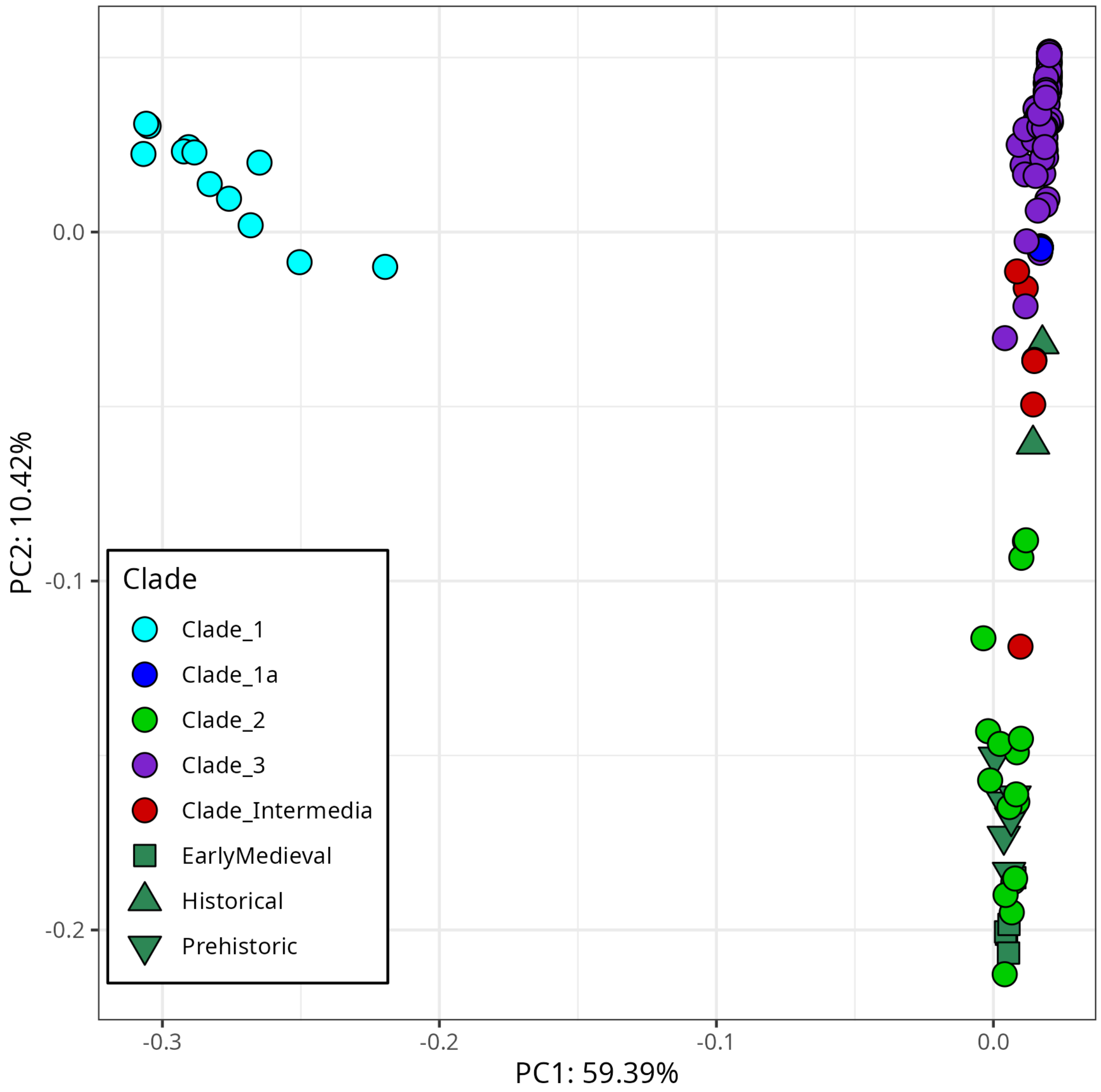


**Supplementary Figure S19.** PCA of 5,953SNPs found across 45 known virulence genes in *E. rhusiopathiae.*

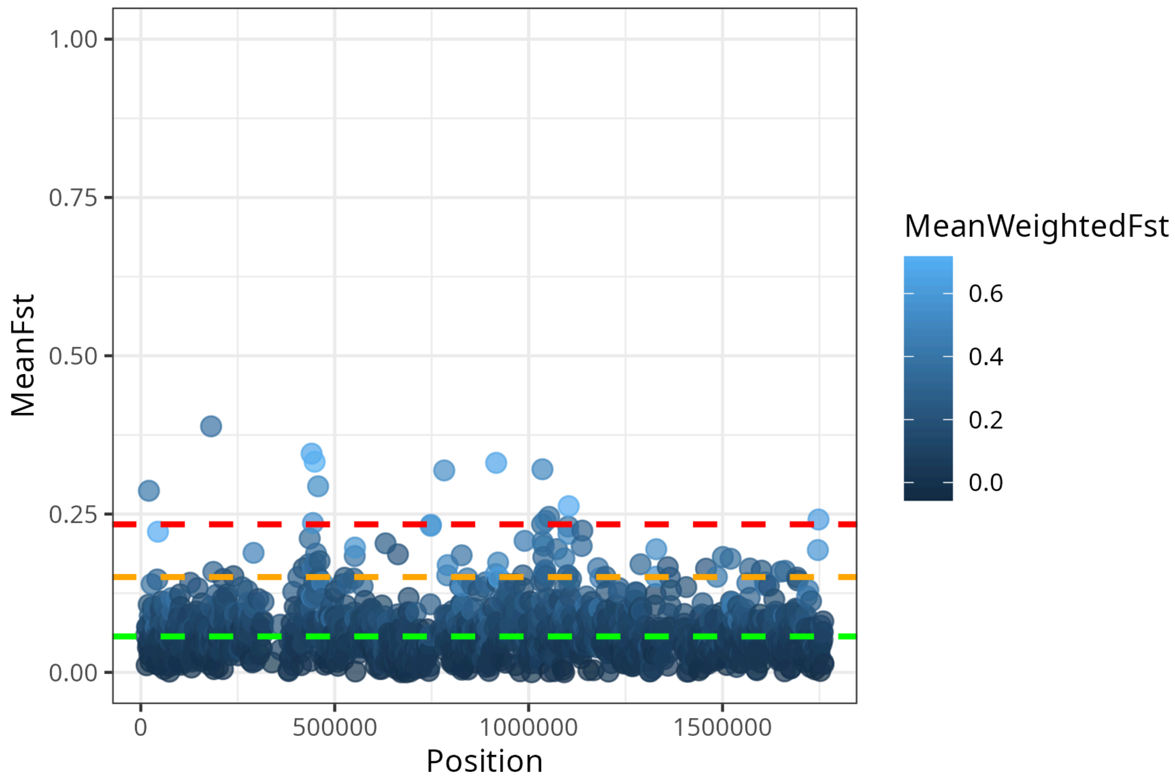


**Supplementary Figure S20.** Wide Genome Weir and Cockerham Fst analysis of Ancient *E. rhusiopathiae* genomes (Medieval and Prehistoric) versus *E. rhusiopathiae* Clade 2 strains.

### Supplementary Tables

**Supplementary Table S1. Laboratory and sequencing statistics.**

**Supplementary Table S2. Pathogen screening results:** Number of validated sequences after KrakenUniq screening and validation by mapping.

| Individual | Hit | Sequences |
|---|---|---|
| GSP013 | *Yersinia pestis* | 4,425 |
| GSP013 | *Erysipelothrix rhusiopathiae* | 4,712 |
| GSP013 | *HBV* | 1 |
| GSP004 | *HBV* | 1 |
| GSP005 | *HBV* | 1 |
| GSP011 | *HBV* | 5 |
| GSP014 | *HBV* | 72 |

**Supplementary Table S3:** Radiocarbon dates from human bone at Grotta della Spinosa

| Lab code | Context (layer, level, quadrant) | Element | 14C determination (BP) | Calibrated date (cal BCE) | Modelled date (cal BCE) |
|---|---|---|---|---|---|
| OxA-18391 | US8, liv. II, G1 | Tibia | 4555±34 | 3370–3110 (1σ)<br>3490–3100 (2σ) | *3220–3100 (1σ)*<br>*3360–3090 (2σ)* |
| Beta-465437 | US8, liv. VI, C2 | Long bone | 4530±30 | 3360–3100 (1σ)<br>3370–3100 (2σ) | *3210–3100 (1σ)*<br>*3350–3090 (2σ)* |
| Beta-456214 | US8, liv. VI, H1 | Humerus | 4520±30 | 3360–3100 (1σ)<br>3360–3100 (2σ) | *3210–3100 (1σ)*<br>*3340–3030 (2σ)* |
| OxA-18392 | US8, liv. VII, H1–I2 | Humerus | 4503±33 | 3340–3100 (1σ)<br>3360–3090 (2σ) | *3200–3090 (1σ)*<br>*3300–3020 (2σ)* |
| Beta-465436 | US8, liv. IV, C2 | Long bone | 4500±30 | 3340–3100 (1σ)<br>3360–3090 (2σ) | *3200–3090 (1σ)*<br>*3300–3030 (2σ)* |
| Beta-456208 | US8, liv. II, G1 | Humerus | 4490±30 | 3340–3100 (1σ)<br>3350–3030 (2σ) | *3230–3090 (1σ)*<br>*3300–3020 (2σ)* |
| Beta-456209 | US8, liv. III, C2 | Tibia | 4470±30 | 3330–3040 (1σ)<br>3340–3020 (2σ) | *3190–3020 (1σ)*<br>*3300–3010 (2σ)* |
| Beta-456207 | US8, liv. II, G1 | Femur | 4450±30 | 3330–3020 (1σ)<br>3340–2930 (2σ) | *3180–3010 (1σ)*<br>*3310–2950 (2σ)* |
| Beta-456212 | US8, liv. IV, H1–H2 | Femur | 4450±30 | 3330–3020 (1σ)<br>3340–2930 (2σ) | *3180–3010 (1σ)*<br>*3320–2950 (2σ)* |
| Beta-456206 | US8, liv. I, I1 | Femur | 4380±30 | 3020–2920 (1σ)<br>3100–2910 (2σ) | *3080–2950 (1σ)*<br>*3100–2920 (2σ)* |
| Beta-456211 | US8, liv. IV, G1 | Tibia | 4380±30 | 3020–2920 (1σ)<br>3100–2910 (2σ) | *3080–2950 (1σ)*<br>*3100–2920 (2σ)* |
| Beta-456213 | US8, liv. V, H1–I1 | Humerus | 4380±30 | 3020–2920 (1σ)<br>3100–2910 (2σ) | *3080–2950 (1σ)*<br>*3100–2920 (2σ)* |
| OxA-18389 | US8, liv. I, F1/F-1 | Rib | 4371±32 | 3020–2920 (1σ)<br>3100–2900 (2σ) | *3080–2940 (1σ)*<br>*3100–2910 (2σ)* |
| Beta-456210 | US8, liv. III, G1 | Ulna | 4370±30 | 3020–2920 (1σ)<br>3100–2900 (2σ) | *3080–2940 (1σ)*<br>*3100–2910 (2σ)* |
| OxA-18390 | US8, liv. I, F1/F-1 | Vertebra | 4364±33 | 3020–2910 (1σ)<br>3100–2900 (2σ) | *3080–2940 (1σ)*<br>*3100–2910 (2σ)* |
| Beta-4562051 | US8, liv. I, C2 | Femur? | 4330±30 | 3010–2890 (1σ)<br>3030–2890 (2σ) | *3020–2950 (1σ)*<br>*3080–2900 (2σ)* |
| Start Boundary | | | *3290–3140 (1σ)*<br>*3380–3120 (2σ)* | | |
| Duration Interval | | | *200–390 (1σ)*<br>*120–490 (2σ)* | | |
| End Boundary | | | *2980–2890 (1σ)*<br>*3010–2860 (2σ)* | | |

**Supplementary Table S4. Isotopic analysis of GSP013/GSP005**: Quality control parameters and isotopic values for the adolescent (sample number GSP5) and mean values for the overall sample from Grotta della Spinosa [2].

| Sample | Collagen yield (wt%) | %C | %N | C:N | δ13CV-PDB (‰) | δ15NAIR (‰) |
|---|---|---|---|---|---|---|
| GSP5 | 2.7 | 29.7 | 11.0 | 3.1 | -19.9 | 9.2 |
| Grotta della Spinosa mean (n=30) | 3.4 | 35.9 | 13.2 | 3.2 | -20.0 | 9.7 |

**Supplementary Table S5.** ***Yersinia pestis*** **phylogenetic and genomics dataset.**

**Supplementary Table S6.** ***Eryspelothrix spp.*** **screening.**

**Supplementary Table S7.** ***Y. pestis*** **and** ***Eryspelothrix*** **co-occurrance.**

**Supplementary Table S8.** ***E. rhusiopathiae*** **phylogenetic and genomics dataset.**

**Supplementary Table S9.** ***E. rhusiopathiae*** **virulence genes.**

**Supplementary Table. S10 High Fst Genes in Ancient versus Clade 2 genomes**

| Start | End | Locus | Gene | Description |
|---|---|---|---|---|
| 20691 | 21102 | NCTC8163_00015 | Predicted protein | hypothetical protein |
| 181127 | 181301 | NCTC8163_00172 | Predicted protein | hypothetical protein |
| 439916 | 440825 | NCTC8163_00417 | Predicted protein | hypothetical protein |
| 443374 | 444307 | NCTC8163_00422 | FrsA | Fermentation-respiration switch esterase |
| 447685 | 448723 | NCTC8163_00426 | Predicted protein | hypothetical protein |
| 456674 | 457655 | NCTC8163_00433 | ArsR | ArsR/SmtB family transcription factor |
| 782090 | 782684 | NCTC8163_00776 | Predicted esterase | alpha/beta hydrolase |
| 915981 | 916767 | NCTC8163_00906 | AspD | L-aspartate dehydrogenase, NAD(P)-dependent |
| 1034190 | 1035231 | NCTC8163_01016 | ElyC | Lipid carrier protein ElyC involved in cell wall biogenesis, DUF218 family |
| 1035325 | 1035862 | NCTC8163_01017 | RimL | Protein N-acetyltransferase, RimJ/RimL family |
| 1043257 | 1043956 | NCTC8163_01028 | CofE | Coenzyme F420-0:L-glutamate ligase |
| 1052194 | 1053301 | NCTC8163_01035 | CamS | Repeat domain of CamS sex pheromone cAM373 precursor and related proteins |
| 1103082 | 1103604 | NCTC8163_01099 | PfpI-like | type 1 glutamine amidotransferase (GATase1)-like domain found in PfpI from Pyrococcus furiosus |
| 1747280 | 1747589 | NCTC8163_01717 | NtpF | Archaeal/vacuolar-type H+-ATPase subunit F/Vma7 |
